# Encyclopaedia of family A DNA polymerases localized in organelles: Evolutionary contribution of bacteria including the proto-mitochondrion

**DOI:** 10.1101/2023.08.28.554543

**Authors:** Ryo Harada, Yoshihisa Hirakawa, Akinori Yabuki, Eunsoo Kim, Euki Yazaki, Ryoma Kamikawa, Kentaro Nakano, Marek Eliáš, Yuji Inagaki

## Abstract

DNA polymerases (DNAPs) synthesize DNA from deoxyribonucleotides in a semi-conservative manner and serve as the core of DNA replication and repair machineries. In eukaryotic cells, there are two genome-containing organelles, mitochondria and plastids, that were derived from an α-proteobacterium and a cyanobacterium, respectively. Except for rare cases of genome-lacking mitochondria and plastids, both organelles must be served by nucleus-encoded DNAPs that localize and work in them to maintain their genomes. The evolution of organellar DNAPs has yet to be fully understood because of two unsettled issues. First, the diversity of organellar DNAPs has not been elucidated in the full spectrum of eukaryotes. Second, it is unclear when the DNAPs that were used originally in the endosymbiotic bacteria giving rise to mitochondria and plastids were discarded, as the organellar DNAPs known to date show no phylogenetic affinity to those of the extant α-proteobacteria or cyanobacteria. In this study, we identified from diverse eukaryotes 134 family A DNAP sequences, which were classified into 10 novel types, and explored their evolutionary origins. The subcellular localizations of selected DNAPs were further examined experimentally. The results presented here suggest that the diversity of organellar DNAPs has been shaped by multiple transfers of the Pol`I` gene from phylogenetically broad bacteria, and their occurrence in eukaryotes was additionally impacted by secondary plastid endosymbioses. Finally, we propose that the last eukaryotic common ancestor may have possessed two mitochondrial DNAPs, POP and a candidate of the direct descendant of the proto-mitochondrial DNAP, rdxPolA, identified in this study.

## Introduction

Multiple endosymbiotic events played pivotal roles in the evolution of eukaryotes. The most ancient endosymbiotic event that is currently recognizable likely occurred between the primordial (proto)eukaryotic or archaeal cell (host) and an α-proteobacterium (endosymbiont). This event gave rise to mitochondria, which operate metabolic pathways that are critical for cell viability, such as oxidative energy production, amino acid synthesis, β-oxidation of fatty acids, and Fe-S cluster assembly, in modern eukaryotic cells (Roger et al. 2017). Another bacterial endosymbiosis took place later than mitochondrial endosymbiosis in eukaryotic evolution and yielded a photosynthetic organelle, the plastid. In primary plastid endosymbiosis, the host and the endosymbiont were the common ancestor of Archaeplastida and a cyanobacterium, respectively (Sibbald and Archibald 2020). The host organism involved in the primary endosymbiosis then diverged into green algae plus land plants (Chloroplastida), glaucophyte algae (Glaucophyta), and red algae (Rhodophyta) plus their non-photosynthetic relatives, so that their plastids are termed primary plastids. Photosynthesis further spread into phylogenetically diverged eukaryotes through multiple “secondary endosymbiosis,” in which a green or red alga was engulfed by a heterotroph and transformed subsequently into a “complex plastid” (Sibbald and Archibald 2020). Green alga-derived complex plastids have been found in two distantly related branches in the tree of eukaryotes, namely Euglenophyceae and Chlorarachniophyta (ignoring a few dinoflagellates also bearing chlorophyte-derived plastids (Kamikawa et al. 2015; Sarai et al. 2020; Matsuo et al. 2022)). On the other hand, the complex plastids in Ochrophyta, Myzozoa, Haptophyta, and Cryptophyceae can be traced back to red algal endosymbionts.

Reflecting the bacterial origins of mitochondria and plastids, the two organelles typically retain their own genomes, which are however highly reduced compared to their free-living relatives (i.e., α-proteobacteria and cyanobacteria). Disregarding sporadic cases related to the insertion of mobile genetic elements into mitochondrial (e.g., Burger et al. 1999; Nishimura et al. 2019) or plastid genomes (e.g., Kim et al. 2015), organellar genomes do not encode proteins involved in DNA replication and repair. Hence, DNA maintenance in organelles entirely relies on the set of proteins encoded in the nuclear genome, synthesized in the cytosol, and then translocated into the corresponding organelle. DNA polymerases (DNAPs), which catalyze the synthesis of the nascent DNA strand by referring to the template strand, are the core components that are central to DNA replication.

Nucleus-encoded organellar DNAPs have been reported from diverse eukaryotes (Klingbeil et al. 2002; Graziewicz et al. 2006; Moriyama et al. 2008; Mukhopadhyay et al. 2009; Moriyama et al. 2011; Moriyama and Sato 2014; Janouškovec et al. 2015; Hirakawa and Watanabe 2019; Harada et al. 2020; Harada and Inagaki 2021). DNA-dependent DNAPs are generally divided into six families (A, B, C, D, X, and Y) (Filée et al. 2002), and organellar DNAPs belong to family A (famA) that is typified by the bacterial DNA polymerase `I` (Pol`I`). Bacterial Pol`I` is responsible for filling the gap between the Okazaki fragments synthesized by Pol`III` (family C) during the lagging strand synthesis (Okazaki 2017). Pol`III` has been retained by the chromatophore, a cyanobacterium-derived photosynthetic organelle endosymbiotically established in the rhizarian genus *Paulinella* independently on the primary plastid (Nowack et al. 2008). In contrast, Pol`III` is absent from mitochondria and plastids, and Pol`I`-related (i.e., family A) DNAPs play a major role in their genome replication (Graziewicz et al. 2006; Parent et al. 2011).

The known organellar DNAPs can be subdivided into five phylogenetically distinct types, namely Polγ, POP, PREX, PolIA, and PolIBCD+ (Harada and Inagaki 2021). POP has been found in phylogenetically diverse eukaryotic lineages and can be targeted to mitochondria, plastids, or both organelles in the same cell (i.e., dual-targeted) depending on the species (Kimura et al. 2002; Christensen et al. 2005; Mori et al. 2005; Ono et al. 2007; Moriyama et al. 2008; Parent et al. 2011; Hirakawa and Watanabe 2019). The origin of POP remains uncertain, as it forms a lineage of its own, lacking any specific relatives in famA DNAPs phylogenies presented so far. In contrast, Polγ, PREX, Pol`I`A, and Pol`I`BCD+ appear to be lineage-specific. Polγ is the mitochondrial DNAP found exclusively in opisthokonts. PREX has been reported only from apicomplexans and closely related lineages (chrompodellids and squirmids), and works in their plastids (Janouškovec et al. 2019). Pol`I`A has been found in all three major classes (Euglenoidea, Diplonemea, and Kinetoplastea) comprising the phylum Euglenozoa, and Pol`I`BCD+ is restricted to Diplonemea and Kinetoplastea (Harada et al. 2020; Harada and Inagaki 2021). The origin of the different organellar DNAPs as elucidated by phylogenetic analyses varies among them. PREX was acquired from a bacterial source, although the specific bacterial group has not been resolved (Janouškovec et al. 2019). Polγ and Pol`I`BCD+ are derived from different bacteriophages (Filée et al. 2002; Harada and Inagaki 2021), and Pol`I`A most likely shares the ancestry with two closely related nuclear famA DNAPs, Polθ and Polν (Harada and Inagaki 2021). Importantly, none of the five types of organellar DNAP has any phylogenetic affinity to the Pol`I` sequences of α-proteobacteria or cyanobacteria, which are the origins of mitochondria and plastids, respectively.

While the recognition of the five aforementioned types of famA DNAP and their taxonomic distribution among eukaryotes established by previous studies defines the mechanistic basis for organellar genome replication and repair in the vast portion of the eukaryote phylogenetic diversity, there are indications they do not constitute a full picture of the organellar DNAP spectrum. Indeed, putative organelle-localized famA DNAPs unrelated to any of the five well-defined types were identified in a single red alga (Moriyama et al. 2008) and a single species belonging to Discoba (Gray et al. 2020). In this study, we conducted a comprehensive survey of famA DNAPs in eukaryotes, which led to the discovery of several novel types of organellar DNAP, including a previously unnoticed DNAP that appears to be a direct genetic legacy of the α-proteobacterial ancestor of the mitochondrion.

## Results & Discussion

### Diversity and subcellular localizations of the novel types of famA DNAP in eukaryotes

We surveyed various sequence databases collectively covering the whole breadth of the eukaryote phylogenetic diversity as captured by sequencing efforts so far. Phylogenetic analyses and sequence curation enabled us to classify the retrieved sequences and discriminate genuine eukaryotic genes from bacterial contaminants. We focused on identifying putative organellar DNAPs in major eukaryote lineages that have not been investigated in this regard in the past and on discovering possible novel famA DNAP types unrelated to POP, Polγ, PREX, Pol`I`A, or Pol`I`BCD+. In this study, we identified 134 famA DNAPs in diverse eukaryotes that were distantly related to any of the previously known eukaryotic famA DNAPs. We prepared a “global famA DNAP” alignment by including diverse bacterial and phage Pol`I` sequences, the six previously known types of eukaryotic famA DNAP sequences, and the famA DNAP sequences identified in this study. From this alignment, we reconstructed the maximum likelihood (ML) tree and calculated ultra-fast bootstrap supports (UFBPs) for bipartitions in the ML tree (Hoang et al. 2018). Based on the global famA DNAP phylogeny (Fig. 1), the novel types of famA DNAP coalesced into ten clades, which we denote “alvPolA,” “abanPolA,” “acPolA,” “rgPolA,” “chlnmPolA,” “eugPolA,” “pyramiPolA,” “chloroPolA,” “cryptoPolA,” and “rdxPolA” (Table 1). The putative domain structure of the representative of each of the ten novel DNAPs is provided in Fig. S1. Importantly, the global famA DNAP phylogeny demonstrated that none of the ten novel DNAPs appeared to be related to any previously known famA DNAPs in eukaryotes (Fig. 1). For each novel famA DNAP type introduced above, we described the phylogenetic distribution and evaluated the subcellular localization primarily by considering experimental data. In the case of no experimental data for a particular famA DNAP, we referred to the results from *in silico* prediction tools (Supplementary Table S1). To complete the atlas of organellar DNAPs, we listed the predicted subcellular localizations of the representative sequences of POP, Polγ, PREX, Pol`I`A, and Pol`I`BCD in Table S1. If experimental evidence for mitochondrial or plastid localization of a particular DNAP exists, we also provided the corresponding reference in Table S1.

**Figure 1.**
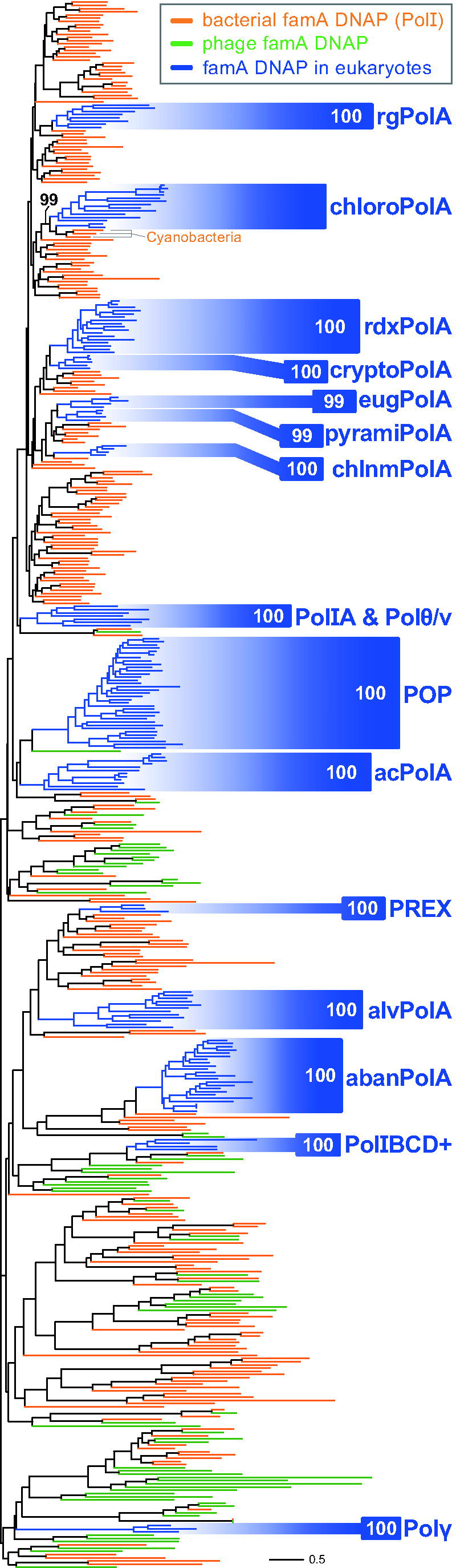
Global phylogeny of family A DNA polymerases (famA DNAPs) sampled from bacteria, phages, and eukaryotes. The ML tree was inferred from an alignment comprising 488 famA DNAP sequences with 355 aa positions. The terminal branches colored in orange, green, and blue represent the famA DNAPs originated from bacteria, phages, and eukaryotes, respectively. The branches of 15 types of famA DNAP found in eukaryotes are shaded in blue. We present the ultra-fast bootstrap support (UFBP) value for the node uniting the paraphyletic chloroPolA sequences and cyanobacterial Pol`I` sequences on the corresponding node. Otherwise, the UFBP value for the monophyly of each type of eukaryotic famA DNAP is shown in the corresponding shaded area.

**Table 1.**
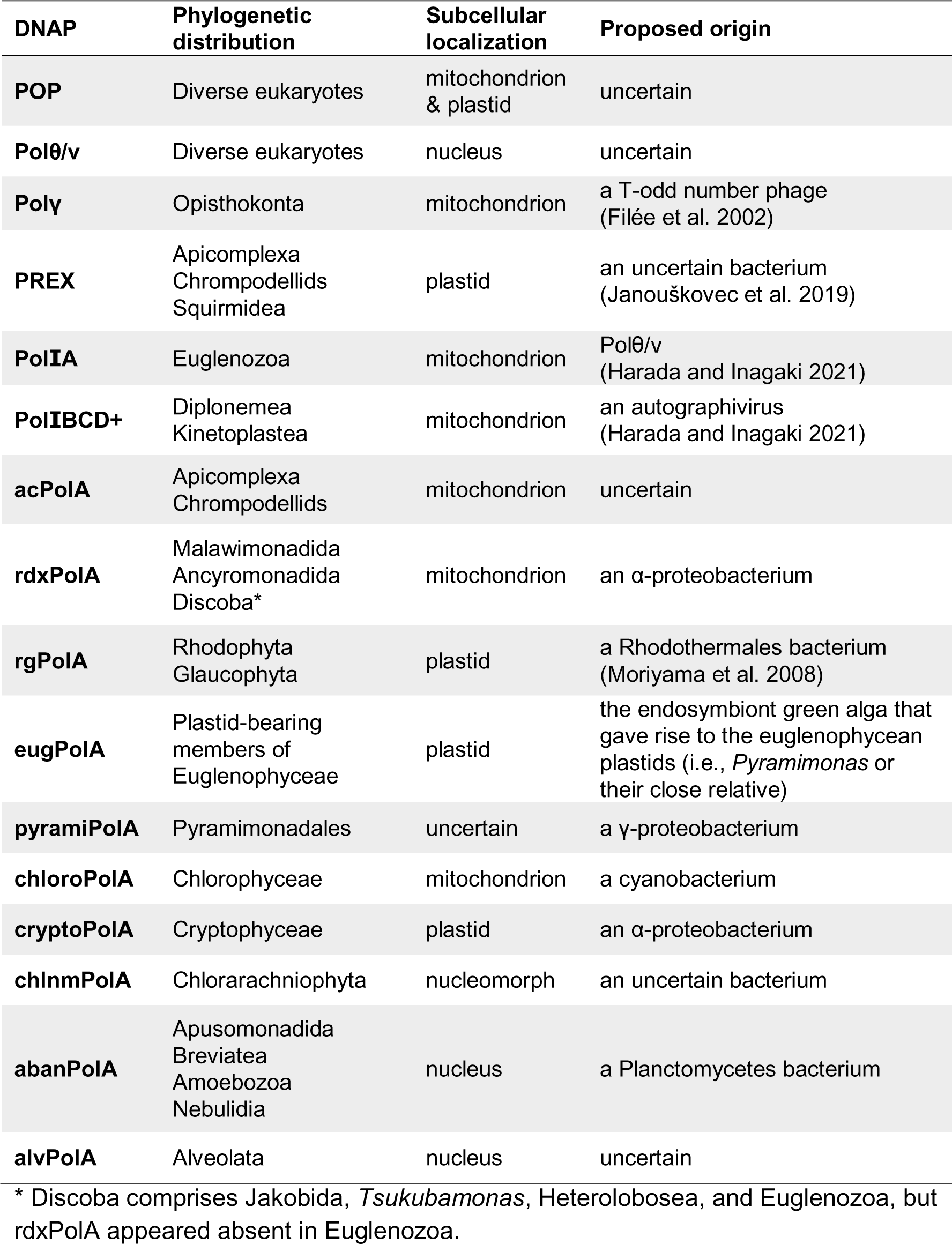
List of family A DNA polymerases (DNAPs) in Eukaryota known to date.

### alvPolA

Our survey revealed that certain representatives of Ciliophora, Dinoflagellata, Squirmidea, Chrompodellida, and Apicomplexa, which all belong to Alveolata, possess previously undescribed famA DNAP sequences that formed a clade supported by an UFBP of 100% in the global famA DNAP phylogeny (Fig. 1). We hence denote the DNAP sequences found in alveolates as “alvPolA.” It is currently ambiguous whether alvPolA is ancestral for Alveolata as a whole, as this particular DNAP type was not found in the sequence data available for Colponemidia, a lineage sister to all other alveolates (Tikhonenkov et al. 2020). Furthermore, this type of DNAP is sporadically found in this group. For instance, the alvPolA sequences were found solely in two heterotricheans in Ciliophora, and only a single gregarine appeared to possess alvPolA among the apicomplexans studied here (Table S1). Given the occurrence of this DNAP type in plastid-lacking members of Alveolata, namely ciliates, it is unlikely to be plastid-localized. Indeed, *in silico* analyses of the N-terminal pre-sequences of alvPolA provided no strong evidence for organellar localization of this DNAP type (Table S1). Combined, we regard alvPolA as nucleus-localized.

### abanPolA

The global famA DNAP phylogeny reconstructed a clade of the novel DNAP sequences identified in the members of three subclades of the eukaryotic “megagroup” Amorphea, namely Apusomonadida, Breviatea, and Amoebozoa, as well as Nebulidia, a subgroup of the supergroup Provora (hereafter designated as “abanPolA”). All the abanPolA-bearing eukaryotes lack a plastid. Furthermore, abanPolA is present in the breviates *Pygsuia biforma* and *Lenisia limosa*, whose mitochondria are highly reduced with no genome (Table S1). Thus, the phylogenetic distribution of abanPolA indirectly but strongly argues for this DNAP type being involved in maintaining the nuclear genome. Consistent with the above notion, little sign of a targeting signal for any organelle was suggested by the *in silico* analyses of the full-length abanPolA sequences (Table S1).

### acPolA

A group of previously unclassified DNAP sequences identified in two chrompodellids and 12 apicomplexans formed a clade with an UFBP of 100%. Prior to this study, the cellular function of the *Plasmodium falciparum* representative was experimentally confirmed (Reesey 2017). The knockdown of the DNAP gene caused a decrease in the amount of mitochondrial DNA, suggesting that this DNAP participates in DNA maintenance in the *P. falciparum* mitochondrion. By extrapolating the result from the experiment on *P. falciparum* to other apicomplexans and chrompodellids, the novel DNAPs discussed here are regarded as mitochondrion-localized, although bioinformatic assessment of this notion is hampered by N-terminal truncation in most of the available acPolA sequences. Recent studies placed a newly recognized parasitic group, Squirmidea, at the base of Apicomplexa and Chrompodellida (Janouškovec et al. 2019; Mathur et al. 2019; Salomaki et al. 2021; Yazaki et al. 2021). We searched for putative mitochondrion-localized famA DNAP sequences in Squirmidea and found only a single candidate, POP, in the transcriptome assembly of *Digyalum oweni*. Thus, the novel famA DNAP group discussed here is most likely specific to the organisms constituting the Apicomplexa + Chrompodellida clade and henceforth termed “acPolA.” In future, the phylogenetic distribution of acPolA should be examined rigorously by surveys of famA DNAPs in high quality transcriptomes of squirmids.

### rgPolA

A type of famA DNAP appeared to be shared among all the eight red algae and two glaucophyte algae surveyed in this study. The ten DNAP sequences, together with the previously known DNAP in the red alga *Cyanidioschyzon merolae* (Moriyama et al. 2008), formed a clade with an UFBP of 100% in the global famA DNAP phylogeny (Fig. 1). We here propose that the novel DNAPs were derived from a single ancestral DNAP. The plastid localization of the *C. merolae* homolog was determined experimentally, suggesting that this type of DNAP is localized in the plastids of red and glaucophyte algae in addition to the dual-targeted POP (Moriyama et al. 2008). We designated this famA DNAP type as “rgPolA” to reflect the restriction to Rhodophyta and Glaucophyta.

### eugPolA, chlnmPolA, and cryptoPolA

We found distinct famA DNAP types that were specific to Euglenophyceae, Chlorarachniophyta, and Cryptophyceae and here termed “eugPolA,” “chlnmPolA,” and “cryptoPolA,” respectively. The global famA DNAP phylogeny united the four eugPolA sequences, four chlnmPolA sequences, and five cryptoPolA into the individual clades supported by UFBPs of 99-100% (Fig. 1).

The eugPolA sequence of the euglenid *Euglena gracilis* was detected specifically in the plastid proteome (Novák Vanclová et al. 2020), suggesting that this DNAP type is plastid-localized. Besides *E. gracilis*, the eugPolA homologs were found in the species belonging to Euglenales and Eutreptiales in Euglenophyceae. *Euglena longa*, which is heterotrophic but retains a non-photosynthetic plastid (Gockel and Hachtel 2000; Füssy et al. 2020), also possesses a eugPolA gene. No eugPolA sequence was detected in the transcriptome of *Rapaza viridis*, which represents a lineage of Euglenophyceae sister to Euglenales and Eutreptiales combined (Yamaguchi et al. 2012; Karnkowska et al. 2023). As *R. viridis* is a kleptoplastidic phagotroph with no permanent plastid, we propose that eugPolA co-occurs with the permanent Pyramimonadales-derived plastids in Euglenophyceae.

In Chlorarachniophyta and Cryptophyceae, four subcellular compartments contain evolutionarily different genomes, i.e., a nucleus, a mitochondrion, a plastid, and a remnant nucleus of endosymbiont (nucleomorph). The subcellular localization of chlnmPolA was investigated by employing the model chlorarachniophyte *Amorphochlora amoebiformis*. As the N-terminal region was absent in the *A. amoebiformis* chlnmPolA sequence reconstructed from the transcriptome data (Table S1), we instead used the N-terminal amino acid residues of the chlnmPolA sequence of the related species *Bigelowiella natans* and expressed it in *A. amoebiformis* as a translational fusion with GFP. This N-terminal leader sequence navigated GFP into the periplastidial compartment (PPC; Figs. 2 and S2). The PPC in chlorarachniophytes harbors a nucleomorph derived from the green algal endosymbiont taken up by the chlorarachniophyte ancestor. Thus, we conclude that chlnmPolA is involved in the DNA maintenance of nucleomorphs in chlorarachniophyte cells.

**Figure 2.**
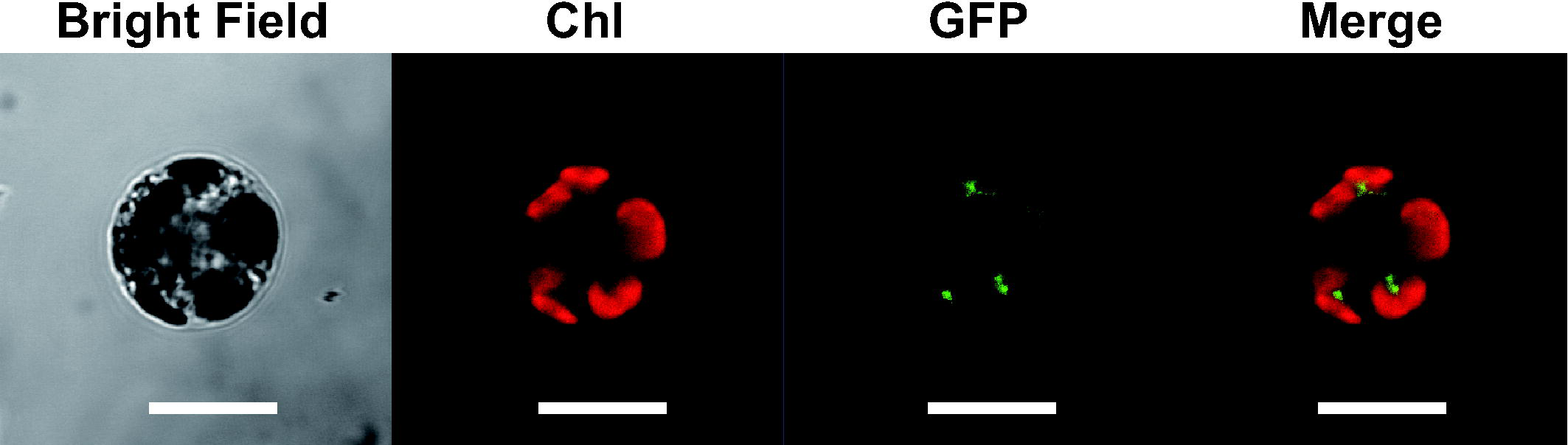
Subcellular localization of GFP fused with the N-terminus of chlnmPolA when expressed in chlorarachniophyte cells. Green fluorescent protein (GFP) fused with the first 100 amino acid residues of the chlorarachniophyte *Bigelowiella natans* chlnmPolA was expressed in the chlorarachniophyte *Amorphochlora amoebiformis*. The red color corresponds to chlorophyll autofluorescence. The green signal represents the GFP localization. The GFP signal was found to be associated tightly with but distinct from plastids, suggesting the protein is localized in the periplastidial compartment. The scale bars are 10 µm.

As no cryptophyte transgene-expression system has been developed, we expressed the N-terminal region of the *Guillardia theta* cryptoPolA as a translation fusion with GFP heterogeneously in the model diatom *Phaeodactylum tricornutum*. This organism was chosen because diatoms and cryptophytes share red alga-derived plastids with the same membrane organization and the plastid protein-targeting mechanism (Nassoury and Morse 2005; Felsner et al. 2011). The green fluorescence signal appeared to overlap with the plastid autofluorescence (Fig. 3A), indicating that the N-terminus of the *G. theta* cryptoPolA was recognized as a plastid targeting signal (PTS) in the diatom cells. This is consistent with the sequence features of N-terminus of the *G. theta* cryptoPolA protein, which includes a predicted signal peptide (Fig. S3A).

**Figure 3.**
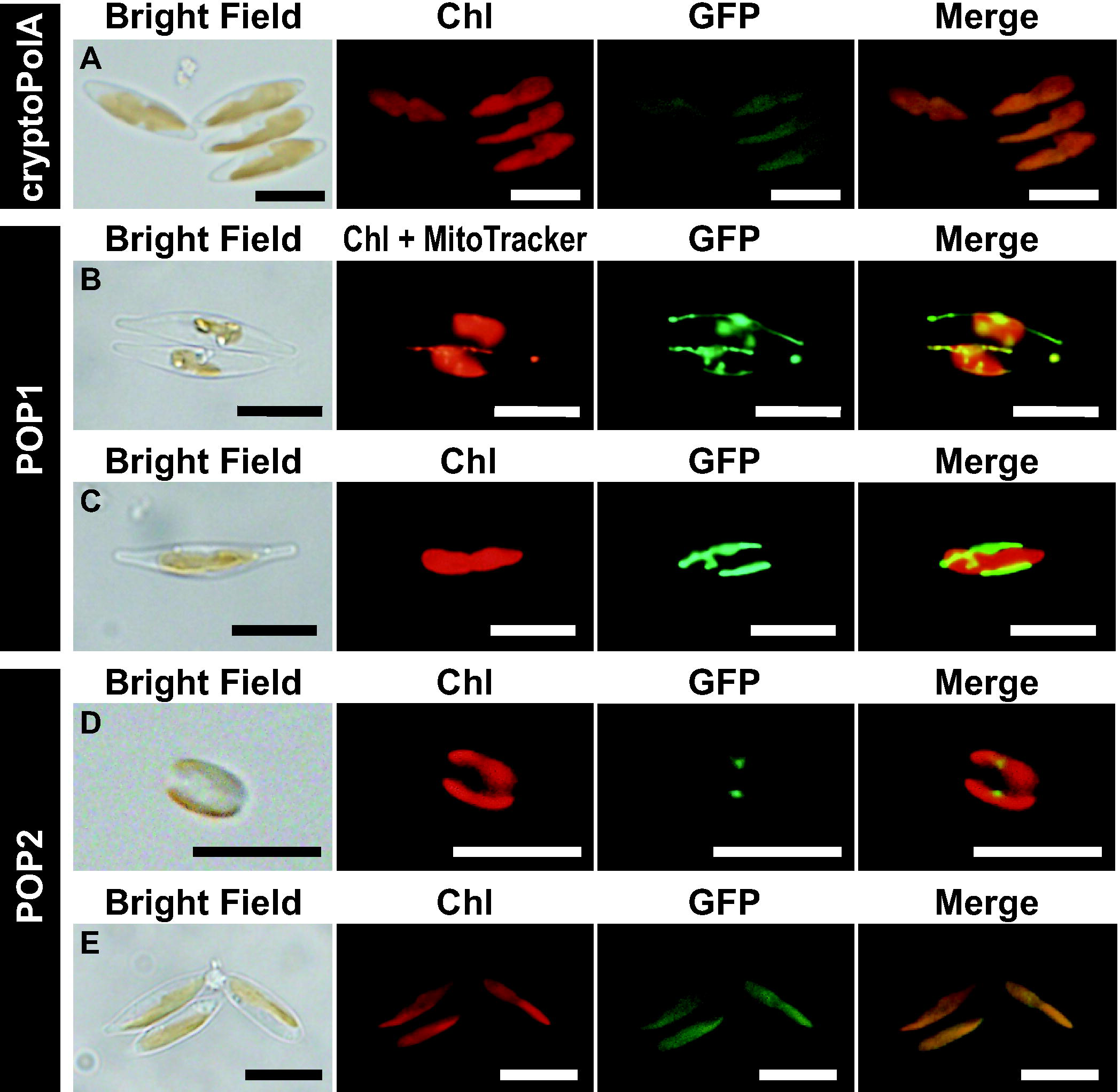
Subcellular localizations of GFP fused with the N-termini of the three organellar family A DNA polymerases found in the cryptophyte *Guillardia theta* when expressed in diatom cells. GFP fused with the N-terminal 332 amino acid residues of *G. theta* cryotoPolA (A), 100 residues of *G. theta* POP1 (B & C), and 516 residues of *G. theta* POP2 (D & E) were heterogeneously expressed in the diatom *Phaeodactylum tricornutum* UTEX642. In A, C, D, and E, the red color corresponds to chlorophyll autofluorescence and the green signal represents GFP. In B, the red color indicates both chlorophyll autofluorescence and mitochondria stained by MitoTracker Orange, and the green signal represents GFP. The scale bars are 10 µm.

Phenylalanine residues appeared to be highly conserved right after the predicted cleavage sites among the plastid-destined proteins in cryptophytes (Gould et al. 2006; Patron and Waller 2007). Intriguingly, in the N-terminus of the *G. theta* cryptoPolA, we found a phenylalanine residue after five amino acids away from the putative cleavage site (Fig. S3A). The same features are conserved in other cryptoPolA proteins for which a complete N-terminal sequence is available (Figs. S3B and S3C), suggesting that cryptoPolA is targeted to the plastid in cryptophytes in general.

Besides cryptoPolA, *G. theta* possesses two distinct POP homologs (Hirakawa and Watanabe 2019), raising the question of their function in the presence of cryptoPolA. Since one of these POPs was a partial sequence and both gene models were potentially inaccurate, we obtained the complete sequences of POP1 (JGI_79451) and POP2 (JGI_199174) from transcriptomic data based on RNA-seq data of PRJNA509798 and PRJNA634446. We repeated the same experiments expressing the POP1 and POP2 N-termini fused to GFP in the diatom cells. In the experiment assessing the localization of POP1, the fluorescence of GFP and MitoTracker Orange, not the plastid autofluorescence, overlapped each other, suggesting that the POP1 N-terminus functioned as a mitochondrial targeting signal (MTS) in the diatom cells (Figs. 3B and 3C). On the other hand, we observed two patterns of the subcellular localization of the green fluorescence signal in the POP2 experiment. In the majority of the transformants, the green fluorescence was observed as small blobs associated with plastids (Fig. 3D). As similar localization patterns have been observed for the PPC proteins of diatoms (Gould et al. 2006; Moog et al. 2020), POP2 may have been localized in the PPC of the diatom cells. In case of POP2 being a genuine PPC protein of *G. theta*, this DNAP needs to be localized in the nucleomorph that is a sole DNA-containing structure in the cryptophyte PPC. Nevertheless, in a small number of the transformants examined, the green fluorescence overlapped with the plastid autofluorescence (Fig. 3E), suggesting that the N-terminus of the *G. theta* POP2 can function as PTS in the diatom cells. However, no typical signal peptide was predicted in the N-terminus of the *G. theta* POP2 *in silico* (Fig.S3D). Although it is difficult to establish the definite subcellular localization of POP2 based on the experiment expressing a cryptophyte protein in a diatom, we propose POP2 as a nucleomorph protein rather than a plastid protein. The subcellular localization of POP2 should be confirmed by electron microscopic observations using an antibody against the DNAP of interest.

### pyramiPolA and chloroPolA

We found two novel types of famA DNAP that are distributed in distinct sublineages of Chloroplastida. First, at least four green algal species of Pyramimonadales possess a unique type of famA DNAP, and these DNAP sequences formed a clade with an UFBP of 99% in the global famA DNAP phylogeny (Fig. 1). The Pyramimonadales-specific famA DNAP is termed as “pyramiPolA.” All the pyamiPolA sequences were predicted to be localized in mitochondria by PredSL (Table S1). Nevertheless, MitoFate and NommPred, the programs designed to detect mitochondrial proteins exclusively, or PredAlgo, a program tuned for green algal proteins, predicted no MTS in any of the four sequences (Table S1). We thus hesitate to discuss further the subcellular localization of pyramiPolA until additional information, preferentially experimental data, becomes available.

Another novel type of famA DNAP was detected in 27 species representing the class Chlorophyceae and belonging to both its principal lineages known as CS and OCC clades. An additional survey of the OneKP database containing transcriptome assemblies of a large phylogenetically diverse set of green algal and plant species (Leebens-Mack et al. 2019) strengthened the case for this novel DNAP type being restricted to Chlorophyceae, and thus we here designate this DNA type as “chloroPolA.” The Chlorophyceae-specific DNAP sequences and cyanobacterial Pol`I` sequences grouped together with an UFBP of 99% in the global famA DNAP phylogeny (Fig. 1). We subjected 11 full-length chloroPolA sequences to *in silico* prediction of subcellular localization, yet the predicted localizations varied depending on the sequence analyzed or the program used (Table S1). We additionally analyzed the same sequences by PredAlgo, a bioinformatic program that was specifically designed to predict protein subcellular localization in green algae (Tardif et al. 2012). Intriguingly, five out of the 11 chloroPolA sequences were predicted as mitochondrion-localized proteins. Thus, we tentatively regard chloroPolA as the DNAP localized in mitochondria in chlorophycean green algae, albeit our conclusion needs to be re-examined experimentally in the future.

### rdxPolA

Gray et al. (2020) reported from the jakobid *Andalucia godoyi* a novel famA DNAP that was predicted mitochondrion-localized and was more closely related to bacterial sequences than to any known previously characterized eukaryotic DNAP type. Our survey of famA DNAP in phylogenetically diverse eukaryotes revealed that specific relatives of the *A. godoyi* DNAP occur not only in other jakobids, but also heteroloboseans and the phylogenetically unique flagellate *Tsukubamonas globosa*, all of which belong to the supergroup Discoba. No putative orthologs were identified in the fourth major Discoba lineage, Euglenozoa, but instead we found them in the two distantly related lineages, Ancyromonadida (three species) and Malawimonadida (two species). All these DNAP sequences formed a clade with an UFBP of 100% in the global famA DNAP phylogeny (Fig. 1). Henceforth, we designate the famA DNAP shared among Malawimonadida, Ancyromonadida, and Discoba as “rdxPolA,” where “rdx” is an abbreviation of the Latin word radix (root). As discussed in the final section, rdxPolA is proposed as the direct descendant of the Pol`I` in the α-proteobacterial endosymbiont that gave rise to the mitochondrion.

We examined the subcellular localization of five rdxPolA proteins (representing all three rdxPolA-possessing Discoba lineages plus a single ancyromonad and a single malawimonad) by expressing their N-terminal regions fused to GFP in the yeast *Saccharomyces cerevisiae*. Significantly, the green fluorescence signal appeared to overlap with mitochondria labeled by MitoTracker Red (Figs. 4A-E), suggesting that the N-termini of rdxPolA proteins were recognized as MTS by the translocon on the mitochondrial membranes in yeast. We anticipate that the subcellular localization deduced from the yeast experiments is applicable to rdxPolA proteins in general. The mitochondrial localization of rdxPolA is independently supported by results of a previous proteomic study on the heterolobosean *Naegleria gruberi* (Horváthová et al. 2021), which lists rdxPolA of this species (XP_002679000.1) as a protein identified in the mitochondrial fraction.

**Figure 4.**
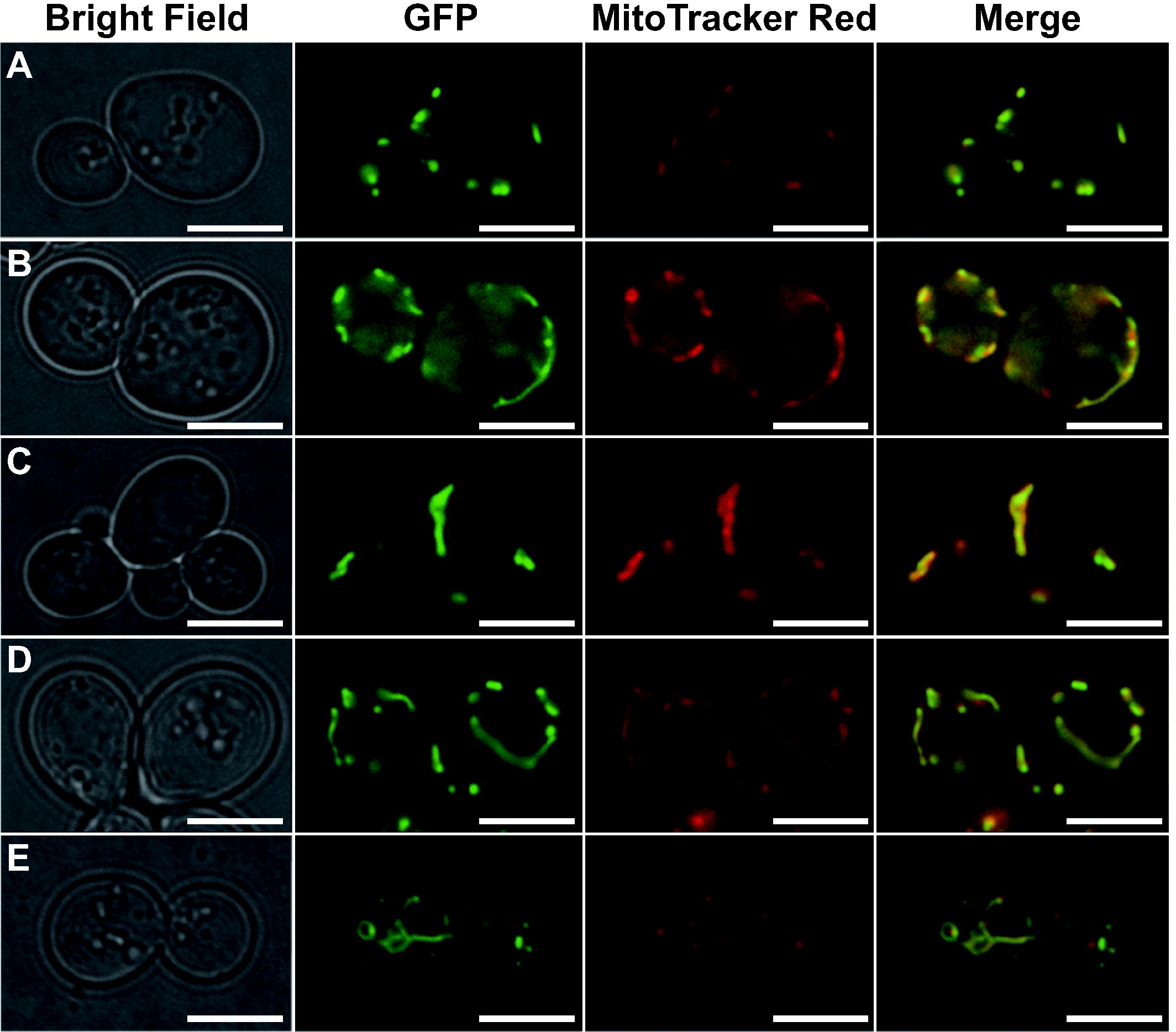
Subcellular localizations of GFP fused with the N-termini of five rdxPolAs when expressed in yeast cells. GFP fused with the N-terminal amino acid residues predicted as the mitochondrial targeting signal (MTS) of rdxPolA proteins from three members of Discoba (*Ophirina amphinema*, *Tsukubamonas globosa*, and *Naegleria gruberi*; A-C), the malawimonad *Gefionella okellyi* (D), and the ancyromonad *Fabomonas tropica* (E). The green signal corresponds to GFP. The red color indicates mitochondria stained by MitoTracker Red. The results presented here demonstrate that the mitochondrial translocons in yeast recognize the N-termini of the five rdxPolAs as the MTS. The scale bars are 5 µm.

### Origins of the novel types of famA DNAP in eukaryotes

The global famA DNAP phylogeny hinted at no direct affinity between any of the eight novel famA DNAP types and phage homologs (Fig. 1). In this section, we explore the origins of the newly defined famA DNAP types in detail (summarized in Table 1). To this end, we prepared eight famA DNAP alignments, each containing one or two novel organellar famA DNAP types, together with a distinct set of bacterial homologs. The individual DNAP alignments were then subjected to both ML and Bayesian phylogenetic analyses. Unfortunately, the origin of three famA DNAP types, acPolA, chlnmPolA, or alvPolA, could not be elucidated with confidence. The monophylies of the three DNAP types were recovered with high statistical support, yet none of the three clades connected to any bacterial Pol`I` sequences with confidence (Figs. S4, S5, and S6). In contrast, as individually discussed below, the origin of the remaining seven novel famA DNAP types could be narrowed down to specific bacterial groups as the most likely donors.

**Figure 5.**
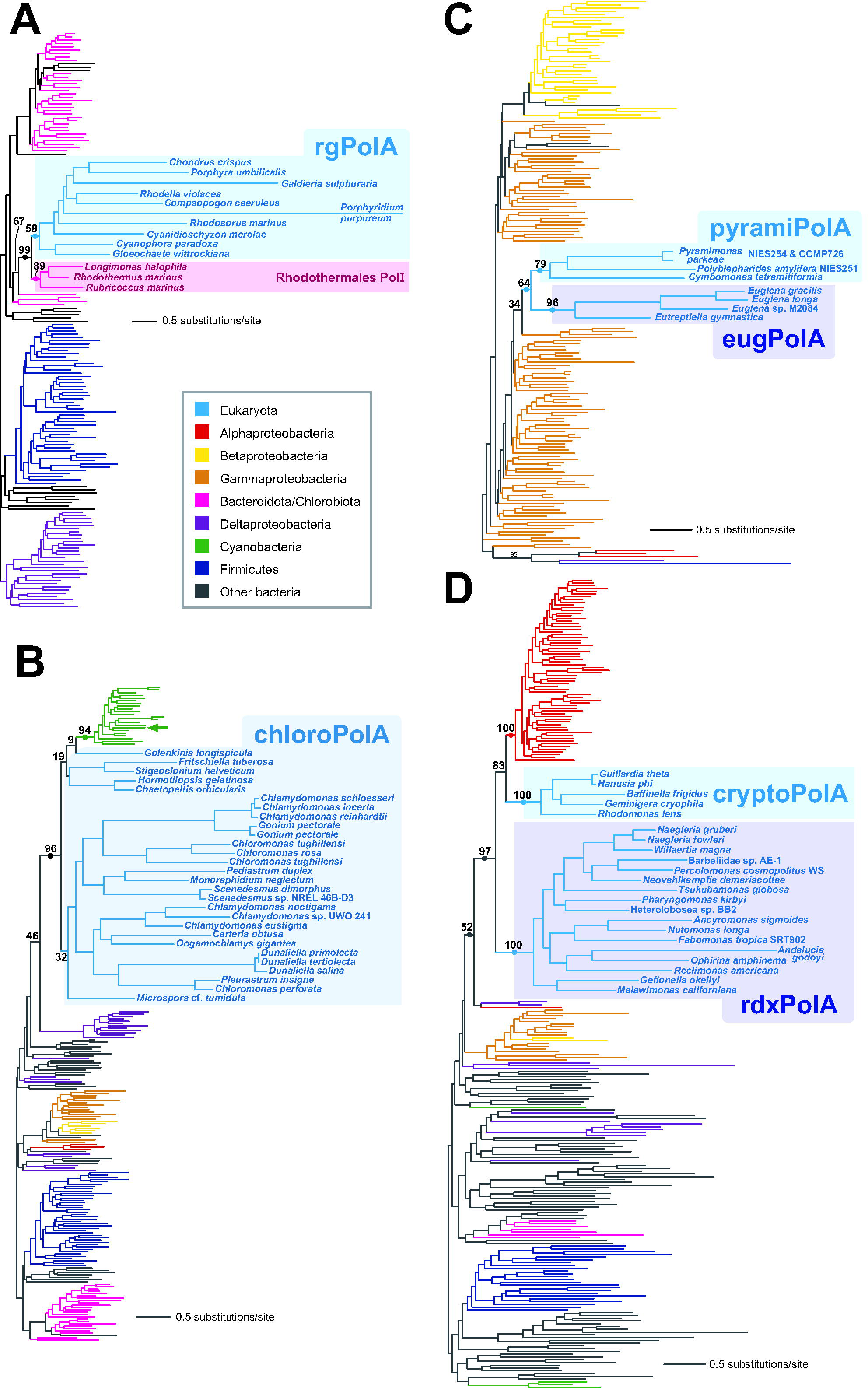
Phylogenies that clarified the origins of six novel types of family A DNA polymerases localized in organelles. The phylogenies assessing the origins of rgPolA (147 sequences, 781 aa positions; panel A), chloroPolA (192 sequences 401 aa positions; panel B), pyramiPolA and eugPolA (161 sequences, 893 aa positions; panel C), and rdxPolA and cryptoPolA (229 sequences, 782 aa positions; panel D). The trees shown here were inferred with the ML phylogenetic method. ML nonparametric bootstrap support values are presented only for nodes key to the origins of the DNA polymerases of interest. In addition, the key nodes that received Bayesian posterior probabilities equal to or greater than 0.95 are marked by dots. The clades/branches are color-coded, as shown in the inset. The Pol`I` of the cyanobacterium *Gloeomargarita lithophora* is marked with a green allow.

#### Planctomycetes origin of abanPolA

Twenty-seven abanPolA sequences formed a clade with an MLBP of 95% and a Bayesian posterior probability (BPP) of 1.0 in the DNAP phylogeny (Fig. S7). The abanPolA clade appeared to be a part of a larger, fully supported clade enclosing the Pol`I` sequences of bacteria belonging to Planctomycetes and those belonging to Myxococcales (part of the traditional paraphyletic δ-proteobacteria). Within this large clade, the abanPolA sequences showed a specific affinity to the Pol`I` sequences of planctomycetes *Novipirellula artificiosorum* and *Tautonia plasticadhaerens* (Fig. S7). Altogether, we conclude that the origin of abanPolA is a laterally transferred Pol`I` gene, and the most plausible candidate for the gene donor is a Planctomycetes bacterium.

#### Rhodothermales origin of rgPolA

Moriyama et al. (2008) demonstrated the specific phylogenetic affinity between the rgPolA homolog in *Cyanidioschyzon merolae* and the Pol`I` sequence of *Rhodothermus marinus*, a bacterium belonging to the phylum Bacteroidota. We here confirmed the origin of rgPolA proposed by Moriyama et al. (2008). In the DNAP phylogeny, the ten rgPolA sequences including the *C. merolae* protein formed a clade with a moderate MLBP, and the rgPolA clade was then grouped with the clade comprising the Pol`I` sequences of *Rhodothermus marinus, Rubricoccus marinus* and *Longimonas halophila*, all of which belong to Rhodothermales in Bacteroidota, with an MLBP of 99% and a BPP of 1.0 (Fig. 5A). The most straightforward interpretation of the DNAP phylogeny is that the origin of rgPolA, which participates in the DNA maintenance in plastids, can be traced back to a Pol`I` gene of a Rhodothermales(-related) bacterium.

#### Cyanobacterial origin of chloroPolA

The chloroPolA sequences, a type of famA DNAP found exclusively in the green algal class Chlorophyceae, and the Pol`I` sequences of cyanobacteria grouped together with an MLBP of 96% and a BPP of 1.0 (Fig. 5B). In this clade, the cyanobacterial Pol`I` sequences formed a subclade with an MLBP of 94% and a BPP of 1.0, whereas the chloroPolA sequences were paraphyletic (Fig. 5B). We here propose that chloroPolA emerged through lateral transfer of a cyanobacterial Pol`I` gene to the common ancestor of Chlorophyceae. In a strict sense, the former proposal demands nesting the chloroPolA clade within the cyanobacterial Pol`I` sequences and thus is inconsistent with the tree topology shown in Fig. 5B. Nevertheless, the large difference in the substitution rate between the cyanobacterial Pol`I` and chloroPolA sequences likely biased tree reconstruction, hindering the recovery of the genuine relationship between chloroPolA and cyanobacterial Pol`I`.

Red and glaucophyte algae possess primary plastids that can be traced back to the “primary endosymbiont,” a cyanobacterium taken up by the common ancestor of Archaeplastida. Nevertheless, the Pol`I` gene, which gave rise to chloroPolA, and that of primary endosymbiont are most likely different. Despite the potential difficulty in elucidating the precise position of the rapidly evolving chloroPolA homologs, the Pol`I` of *Gloeomargarita lithophora*, which has been proposed to be the closest relative of the primary endosymbiont among the cyanobacteria known to date (Ponce-Toledo et al. 2017), was distantly related to the chloroPolA in the DNAP phylogeny (green arrow in Fig. 5B). Moreover, chloroPolA showed a highly restricted distribution within Archaeplastida. Thus, if chloroPolA is assumed as the direct descendant of the Pol`I` in primary endosymbiont, a potentially large number of secondary losses of chloroPolA needs to be invoked after the diversification of the extant members of Archaeplastida. Altogether, the aforementioned hypothesis on the emergence of chloroPolA by lateral gene transfer (LGT) of a Pol`I` gene from a cyanobacterial donor into the common Chlorophyceae ancestor appears more realistic.

#### eugPolA and pyramiPolA: γ-proteobacterial origin and endosymbiotic connection

The eugPolA and pyramiPolA sequences, of which a sister relationship was recovered in the global famA DNAP phylogeny (UFBP of 86%; Fig. 1), were separately subjected to BLAST surveys against the bacterial sequences and matched mainly with the Pol`I` sequences of diverse γ-proteobacteria. Thus, a single alignment containing both eugPolA and pyramiPolA sequences was prepared and subjected to ML phylogenetic analyses. In the resultant ML tree (Fig. 5C), the eugPolA and pyramiPolA sequences grouped together with an MLBP of 64% and a BPP of 0.99, and were placed within the radiation of the γ-proteobacterial Pol`I` sequences. The current plastids in euglenophycean algae descended from a plastid of a green alga belonging to the order Pyramimonadales via endosymbiosis or kleptoplasty (Turmel et al. 2009; Jackson et al. 2018; Karnkowska et al. 2023). Combining the phylogenetic affinity between eugPolA and pyramiPolA (Fig. 5C) with the pyramimonadalean origin of euglenophycean plastids, we here propose that the common ancestor of the extant photosynthetic euglenophyceans acquired the gene encoding pyramiPolA from the endosymbiont and then modified it into a plastid DNAP that corresponds to the ancestral eugPolA. As the γ-proteobacterial Pol`I` sequences enclosed both eugPolA and pyramiPolA sequences in the DNAP phylogeny (Fig. 5C), we here additionally propose the γ-proteobacterial Pol`I` origin of pyramiPolA (and indirectly of eugPolA). We are currently uncertain about the subcellular localization of pyramiPolA (see above). However, this uncertainty is unlikely to spoil the aforementioned hypothesis, as the ancestor of photosynthetic euglenophyceans may have changed the subcellular localization of the endosymbiotically acquired DNAP. It is still possible, though less parsimonious, that the origin of eugPolA is unrelated to the DNAP of the pyramimonadalean endosymbiont that was transformed into the plastid. If so, two distinct γ-proteobacteria donated their Pol`I` genes separately to the common ancestor of the extant photosynthetic euglenophyceans and the common ancestor of Pyramimonadales.

#### cryptoPolA and rdxPolA originated from separate α-proteobacterial PolI genes

The cryptoPolA and rdxPolA sequences appeared to share a phylogenetic affinity to the Pol`I` sequences of α-proteobacteria in BLAST surveys. Hence, we prepared and analyzed a single alignment including both rdxPolA and cryptoPolA. Significantly, the rdxPolA and cryptoPolA sequences grouped together with the α-proteobacterial Pol`I` sequences with an MLBP of 97% and a BPP of 1.0 to the exclusion of the Pol`I` sequences of other bacteria (Fig. 5D). Within this clade, the rdxPolA, cryptoPolA, and α-proteobacterial Pol`I` sequences coalesced into three subclades with full statistical support, and the sister relationship between the cryptoPolA and α-proteobacterial Pol`I` subclades was recovered with an MLBP of 83% and a BPP of 0.67 (Fig. 5D). In the phylogenetic tree of eukaryotes, cryptophytes are included in Cryptista, which further forms the “CAM” clade with *Microheliella maris* and Archaeplastida (Yazaki et al. 2022), and distantly related to any of the rdxPolA-bearing lineages. Thus, if we assume that cryptoPolA and rdxPolA share a common ancestry, such hypothetical famA DNAP may have been lost secondarily in potentially a large number of the lineages that intervene Cryptophyceae, Malawimonadida, Ancyromonadida, and Discoba in the tree of eukaryotes. Notably, cryptoPolA type is present in multiple distantly related members of Cryptophyceae but is also missing from the Goniomonadaceae, the immediate non-photosynthetic (plastid-lacking) sister lineage of Cryptophyceae represented by *Goniomonas* spp. and *Hemiarma marina* (Yazaki et al. 2022). It thus seems that cryptoPolA was established in the common ancestor of Cryptophyceae and its emergence coincides with the establishment of the secondary plastid in the evolution of the Cryptista lineage, in agreement with the plastidial localization of this DNAP (Fig. 3A). This provides an extra argument for the separate origins of cryptoPolA and rdxPolA. Thus, although the mutual relationship of cryptoPolA and rdxPolA cannot be settled with the present data, the DNAP phylogeny can be interpreted as cryptoPolA and rdxPolA being traced back to distinct α-proteobacterial DNAPs.

### Evolution of organellar famA DNAPs in eukaryotes

This study expanded our knowledge of famA DNAPs in eukaryotes. From now on, we will focus on the evolution of organellar famA DNAPs and thus omit the two newly discovered putative nuclear DNAPs, alvPolA and abanPolA, from the discussion below. As summarized in Fig. 6, 16 types of famA DNAP (three types of POP with distinct subcellular localizations counted as one) have been identified in phylogenetically diverse eukaryotes, with 12 of them known or predicted to function in organelles. In this section, we combine the updated diversity of organellar famA DNAPs with the current view on the global eukaryotic phylogeny to paint an integrated picture of the evolutionary history of organellar DNAPs in eukaryotes, dividing the discussion according to the major segments of the eukaryote phylogenetic tree.

**Figure 6.**
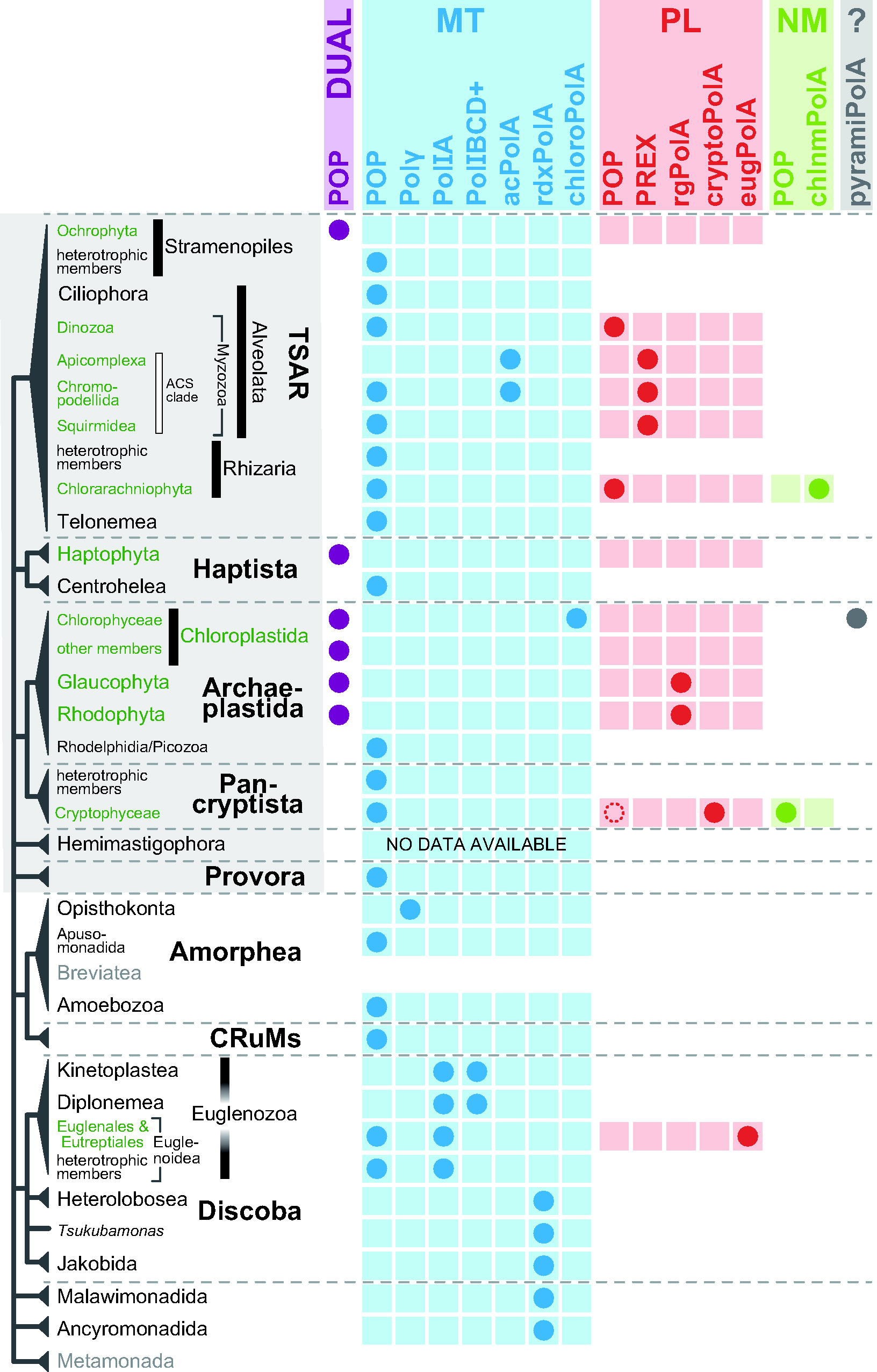
Summary of organellar family A DNA polymerases and their phylogenetic distributions. This figure summarizes the diversity of organellar family A DNA polymerases (DNAPs) and their phylogenetic distributions over the tree of eukaryotes (schematically shown on the left). The presence of each type of organellar DNAP is indicated by a dot colored in blue (mitochondrion-localized; MT), red (plastid-localized; PL), or green (nucleomorph-localized; NM), respectively. pyramiPolA found in green algae of the order Pyramimonadales is shown in a gray dot, as its exact organellar (mitochondrial or plastid) localization remains inconclusive. None of the currently known members of Breviatea or Metamonada, which bear highly reduced mitochondria whose own genomes had been discarded completely, is anticipated to retain any organellar DNAP, and the corresponding rows were left blank. Although we are aware of the laterally acquired POP in a subgroup of choanoflagellates (see Fig. S8), we regard the choanoflagellate POP as an exception and omitted it from this figure. We found no candidate for mitochondrial DNAP in the transcriptomes of hemimastigophorans and thus commented so (“NO DATA AVAILABLE”).

### Mitochondria in Diaphoretickes

Diaphoretickes is a designation of a large-scale taxonomic assemblage (“megagroup”) comprised of Archaeplastida (including Rhodelphidia and Picozoa), Pancryptista, Haptista, Telonemea, and SAR (Adl et al. 2019), recently shown to embrace also Hemimastigophora and Provora (Lax et al. 2018; Tice et al. 2021; Tikhonenkov et al. 2022). We detected POP proteins with predicted mitochondrial localization in representatives of several deeply diverged Diaphoretickes taxa where the nature of the mitochondrial DNAP had not been defined before, including Telonemea, Centrohelea, Microhelida, Rhodelphidia, Picozoa, and Provora (Fig. 6; Table S1). Taken together, all members of Diaphoretickes except Apicomplexa appeared to utilize POPs as the principal DNAP type for DNA replication/repair in mitochondria (some plastid-bearing lineages use a single POP that localizes both mitochondria and plastids; see below for the details) (Fig. 6). Thus, we predict that the common ancestor of Diaphoretickes used POP for DNA maintenance in mitochondria.

In apicomplexan mitochondria, acPolA, instead of POP (completely missing from the group), most likely works as the principal DNAP in their mitochondria (Fig. 6). Some of us previously demonstrated that chrompodellids, the closest relatives of Apicomplexa, possess a mitochondrion-localized POP (Hirakawa and Watanabe 2019), albeit acPolA was additionally identified in chrompodellids in this study (Fig. 6). Intriguingly, as mentioned above our survey detected a mitochondrion-localized POP but provided no evidence for acPolA in Squirmidea, a lineage sister to Apicomplexa and Chrompodellida combined. We thus infer that in the mitochondrion of the common ancestor of the clade of Apicomplexa, Chrompodellida, and Squirmidea (ACS clade; Mathur et al. 2023), like other alveolates including their sister group, Dinozoa, DNA maintenance used to be governed by POP. A novel type of mitochondrial DNAP, acPolA, was then introduced in the common ancestor of Apicomplexa and Chrompodellida, with chrompodellids keeping POP along with acPolA. Nevertheless, the secondary loss of POP having occurred on the branch leading to Apicomplexa suggests acPolA could fully take over the ancestral role of POP in this group.

We revealed that chlorophycean green algae share a unique type of a putative mitochondrion-localized famA DNAP, chloroPolA (Fig. 6). It is possible that another unique mitochondrial famA DNAP, pyramiPolA, was independently introduced to Pyramimonadales, although the bioinformatic evidence for the specific organellar localization is less clear in this case (Supplementary Table S1). Although experimental confirmation is required, chloroPolA, together with POP, is likely involved in DNA maintenance in chlorophycean mitochondria, and it cannot be ruled out that a similar duality exists in Pyramimonadales. Interestingly, the chloroPolA gene in *Chlamydomonas reinhardtii* (Cre17.g736150) was found to be expressed specifically in the zygote, with very limited expression detected in vegetative cells (Joo et al. 2017), pointing to functional specialization of this DNAP.

Two distinct types of DNAP thus appeared to be localized in the mitochondria of some eukaryotes (e.g., POP and acPolA in the chrompodellid mitochondria). Unfortunately, the sequence data we analyzed here are insufficient to conclude whether the two co-existing DNAPs are functionally differentiated or redundant in the mitochondrial DNA maintenance. To address this issue appropriately, the function of each DNAP of interest needs to be examined experimentally.

### Plastids in Diaphoretickes

Diaphoretickes contains multiple plastid-bearing lineages and the repertories of the DNAPs involved in plastid DNA maintenance vary in lineage-specific manners. Chloroplastids, ochrophytes (plastid-bearing species in Stramenopiles), and haptophytes all seem to operate “dual-target POPs,” which can manage DNA maintenance in both mitochondria and plastids (Kimura et al. 2002; Christensen et al. 2005; Mori et al. 2005; Ono et al. 2007; Parent et al. 2011; Hirakawa and Watanabe 2019) (Fig. 6). The red alga *Cyanidioschyzon merolae* was also demonstrated to have a dual-targeted POP, yet supplemented with a plastid-localized rgPolA (Moriyama et al. 2008). Our study has expanded the phylogenetic distribution of the pair of the (presumably) dual-targeted POP and the plastid-localized rgPolA from a single red alga to multiple red algae as well as glaucophytes (Fig. 6). The organismal phylogenies inferred from nucleus-encoded proteins constantly united Chloroplastida and Glaucophyta (Tice et al. 2021; Irisarri et al. 2022; Yazaki et al. 2022), while those inferred from plastid proteins often recovered the sister relationship between Chloroplastida and Rhodophyta (Rodríguez-Ezpeleta et al. 2005; Mackiewicz and Gagat 2014; Figueroa-Martinez et al. 2019). Regardless of the two competing scenarios for the phylogeny of Archaeplastida, the common ancestor of this group may have possessed both dual-targeted POP and plastid-localized rgPolA. The latter was lost not only in Rhodelphidia (retaining a genome-less plastid (Gawryluk et al. 2019)) and Picozoa (that lost the plastid completely (Schön et al. 2021)) but also Chloroplastida. In the case of both POP and rgPolA being localized in a plastid, we need experimental data to clarify the precise functions of the two DNAPs in the plastid DNA maintenance.

The situation concerning plastid DNAPs is specific also in Myzozoa, the alveolate clade combining Dinozoa and ACS clade (Fig. 6). Those dinozoans that have retained a plastid genome (i.e., “core” dinoflagellates) employ plastid-targeted POPs that are distinct from mitochondrion-targeted POPs (Hirakawa and Watanabe 2019). In contrast, the members of ACS clade, i.e., apicomplexans, chrompodellids, and squirmids, exhibit PREX for DNA maintenance in their plastids (Hirakawa and Watanabe 2019; this study), whereas POPs in chrompodellids and squirmids were predicted to be mitochondrion-localized bioinformatically (Fig. 6). At this moment, we are uncertain what type (or types) of DNAP worked in the plastid of the ancestral myzozoan. If the ancestral myzozoan used a single type of a plastid DNAP, either a switch from POP to PREX or that from PREX to POP occurred after the separation of Dinozoa and ACS clade. Alternatively, the ancestral myzozoan may have had both POP and PREX for plastid DNA maintenance. In the latter scenario, differential losses of one of the two types of DNAP need to be invoked on the branches leading to Dinozoa and ACS clade.

Chlorarachniophytes operate distinct POPs localized in the mitochondrion and the plastid (Hirakawa and Watanabe 2019). In addition, we identified an additional famA DNAP, chlnmPolA, which most likely participates in DNA maintenance in the nucleomorph. Besides chlnmPolA, the chlorarachniophyte *Bigelowiella natans* was reported to use an unrelated (family B) DNAP of viral origins for the replication of the nucleomorph genome, in addition to a homolog of DNA polymerase eta (family Y DNAP) presumably involved in nucleomorph genome repair (Suzuki et al. 2016). The ancestral chlorarachniophyte may thus have modified the original green alga-derived machinery for DNA maintenance in the nucleomorph by integrating components with exogenous origins.

The members of another algal nucleomorph-bearing lineage, Cryptophyceae, possess a similar repertory of famA DNAPs to that in chlorarachniophytes. *G. theta* as the reference cryptophyte species specifically has two POPs plus a lineage-specific famA DNAP (cryptoPolA) (Fig. 6). Considering the fluorescent protein tagging in diatom cells, *G. theta* likely uses one of the two types of POP (i.e., POP1) and the lineage-specific famA DNAP (i.e., cryptoPolA) for the DNA maintenance in the mitochondrion and plastid, respectively. Interestingly, the nucleomorph is the best estimate for the subcellular localization of *G. theta* POP2 (see above). Notably, except for the closely related species *Hanusia phi*, our investigation of the genome and/or transcriptome assemblies available for other cryptophyte species did not indicate the presence of the second POP paralog as in *G. theta*. At face value, the restricted distribution of POP2 among cryptophyte algae suggests the recruitment of a POP polymerase for a nucleomorph function only after the divergence of this algal group. However, an alternative interpretation exists: the POP2 transcripts may have been overlooked in at least some of the other cryptophytes. In general, eukaryotic genomes are replicated by family B DNA polymerases (Guilliam and Yeeles 2020), and the cryptophyte nucleomorph genome effectively is a reduced and divergent eukaryotic (specifically red algal) genome. Thus, we speculate that analogously to the presumed role of chlnmPolA in the chlorarachniophyte nucleomorph, the *G. theta* POP2 protein plays an auxiliary role in maintaining the nucleomorph genome in this species (e.g., DNA repair).

### Amorphea plus CRuMs

A large taxonomic assemblage, Amorphea, comprises Opisthokonta, Apusomonadida, Breviatea, and Amoebozoa. Recent phylogenomic studies united collodictyonids, rigifilids, and mantamonads into a clade called CRuMs with high statistical support and further nominated this assemblage as a candidate for the sister branch of Amorphea (Brown et al. 2018). A putative mitochondrion-localized POP has been previously detected in Apusomonadida and Amoebozoa (Hirakawa and Watanabe 2019), and we now found the same for CRuMs. Opisthokonts have been known to use Polγ for DNA maintenance in their mitochondria (Fig. 6), but we have noticed an interesting exception in a subgroup of choanoflagellates (the family Acanthoecidae), which seems to have lost Polγ and instead encodes POP proteins, acquired laterally from a prymnesiophycean haptophyte (Fig. S8). Breviates possess severely modified mitochondria (mitochondrion-related organelles) adapted to the microaerophilic/anaerobic lifestyle and all probably lack a genome (confirmed for the most extensively studied representative *Pygsuia biforma*; Stairs et al. 2014), so it is no surprise that no mitochondrial DNAP candidate was found in this group. By mapping the types of mitochondrial DNAP on the tree of Amorphea, we propose that the common ancestor of this taxonomic assemblage possessed a mitochondrion-localized POP but secondary losses of POP occurred in the two descendant lineages, Opisthokonta and Breviatea. The ancestral opisthokont had discarded the pre-existing POP after the emergence of a lineage-specific mitochondrial DNAP, Polγ. Breviates simply lost the POP due to no use for any mitochondrion-localized DNAP. If CRuMs and Amorphea are truly the closest relatives among eukaryotes, their common ancestor had likely used POP for mitochondrial DNA maintenance.

### Ancyromonadida, Discoba, Malawimonadida, and LECA

Prior to this study, the information on mitochondrial DNAPs in Discoba has been available only for euglenozoans and a single jakobid, *Andalucia godoyi*. In euglenozoans, three types of mitochondrion-localized DNAP, namely POP, Pol`I`A, and Pol`I`BCD+, have been identified (Klingbeil et al. 2002; Harada et al. 2020; Harada and Inagaki 2021), and *A. godoyi* was proposed to use a novel type of famA DNAP for the mitochondrion (Gray et al. 2020). The current study revealed that all the discobids except euglenozoans possess rdxPolA, typified by the *A. godoyi* protein, for mitochondrial DNA maintenance. In the phylogenetic tree of Discoba, jakobids and then *Tsukubamonas globosa* were sequentially separated prior to the split of heteroloboseans and euglenozoans. Thus, we regard that rdxPolA was present prior to the diversification of Discoba as the mitochondrial DNAP and a switch from rdxPolA to POP or Pol`I`A/Pol`I`BCD+ occurred on the branch leading to euglenozoans. The introduction of eugPolA in the ancestor of photosynthetic members of Euglenophyceae should have coincided with the plastid acquisition through green algal endosymbiosis (see above).

Significantly, the origin of rdxPolA may be much earlier than the emergence and diversification of Discoba, as this DNAP is shared by ancyromonads and malawimonads. To elucidate the origin and evolution of rdxPolA, understanding the phylogenetic relationship among the three rdxPolA-bearing lineages is crucial. The situation would be simplest if Malawimonadida, Ancyromonadida, and Discoba constituted a monophyletic group among eukaryotes. However, none of the phylogenomic studies published previously has ever recovered their monophyly (e.g., Tice et al. 2021; Tikhonenkov et al. 2022; Yazaki et al. 2022). In the strict sense, the unrooted phylogenetic analyses are inappropriate to assess whether the rdxPolA-bearing lineages are genuinely monophyletic. However, the lineages of our interest need to form a clan (i.e., be at least potentially monophyletic in the unrooted phylogeny) if they are to form a genuine clade of eukaryotes. We examined the abovementioned issue by analyzing a phylogenomic alignment comprising 340 proteins with no outgroup taxon (Figs. 7A & 7B). As anticipated from the previously published works, the resultant unrooted tree of eukaryotes failed to recover the monophyly of the rdxPolA-bearing lineages (with or without Metamonada) or that of POP-bearing assemblages. In the ML tree, the branch connecting Diaphoretickes and Amorphea plus CRuMs was interrupted by two distinct branches, one uniting Discoba and Metamonada and the other uniting Malawimonadida and Ancyromonadida (Fig. 7A). We subjected the ML tree and eight alternative trees to an approximately unbiased (AU) test (Shimodaira 2002). Significantly, the AU test rejected all the test trees in which the rdxPolA-bearing lineages were forced to be monophyletic regardless of the inclusion or exclusion of Metamonada of which mitochondrion-related organelles lack the genomes, as well as their replication machinery including DNAPs, at a 1% α-level (Trees 7-9; Fig. 7B). Thus, we disfavor the monophyly of the rdxPolA-bearing lineages. As discussed below, this conclusion has important implications for reconstructing details of mitochondrial DNA maintenance in the earliest phases of eukaryote evolution.

**Figure 7.**
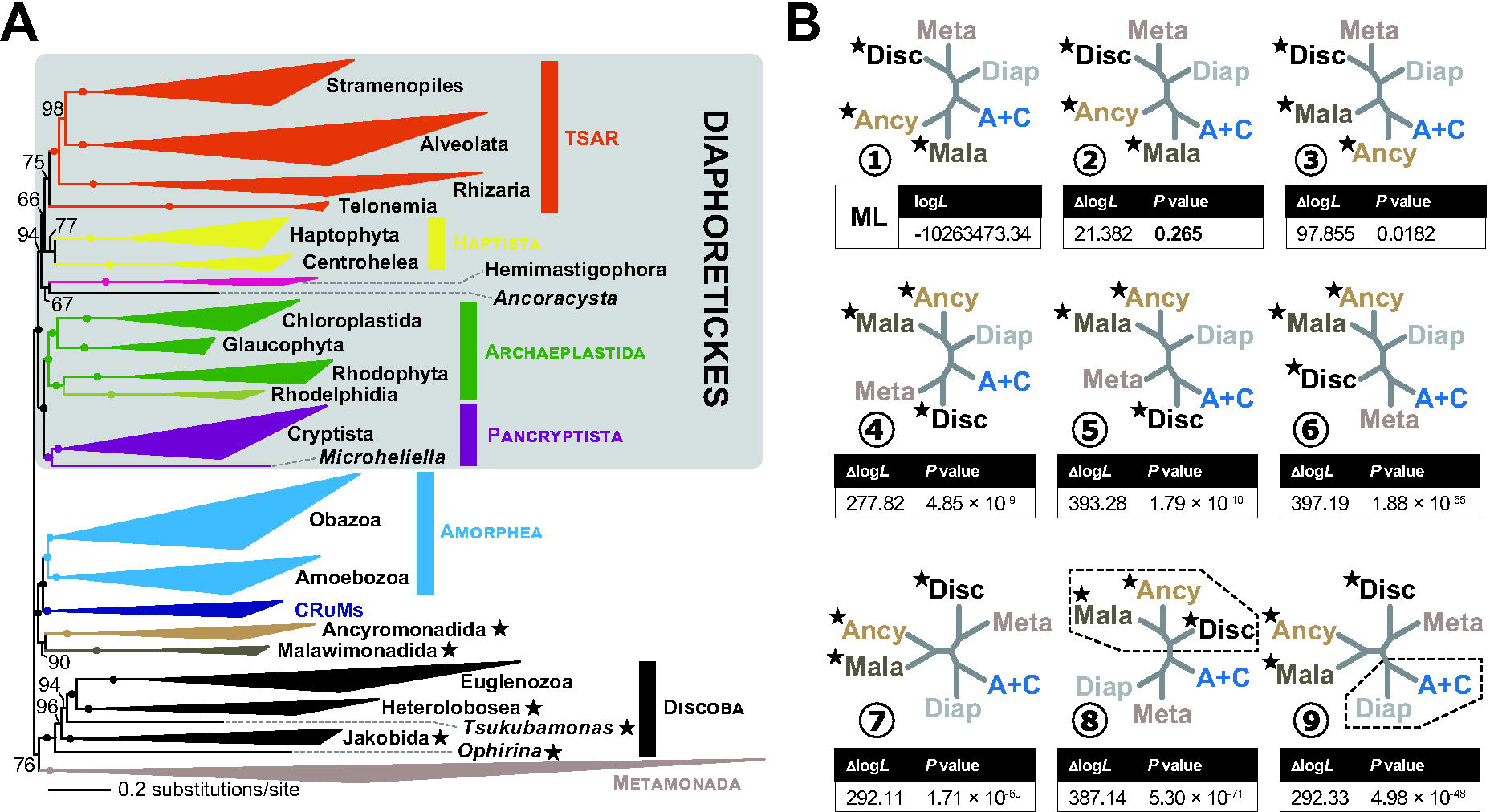
Phylogenomic analyses of eukaryotes. (A) The phylogenetic relationship among 97 eukaryotes was inferred from a phylogenomic alignment comprising 340 proteins by using the ML method. Major clades are shown by triangles. Dots on nodes indicate that the corresponding bipartitions receive ultra-fast bootstrap support (UFBP) values of 100%. The assemblage/lineages/species that use rdxPolA as mitochondrion-localized DNA polymerase are marked by stars. (B) Alternative topologies of the eukaryotic phylogeny assessed in an approximately unbiased test. We here examined only the phylogenetic relationship among six major clades of eukaryotes, namely Diaphoretickes (Diap), Amorphea plus CRuMs (A+C), Ancyromonadida (Ancy), Malawimonadida (Mala), Discoba (Disc), and Metamonada (Meta). Trees 2-7 were almost identical to the ML tree topology (Tree 1), except for the position of Amorphea plus CRuMs. The internal relationship in each major clade (see above) was re-optimized. Tree 8 and 9 are the best trees searched under the topological constraints assuming the monophyly of the POP-bearing and the rdxPolA-bearing lineages, respectively. The dotted lines in Trees 8 and 9 represent the constraints during the tree search described above. The rdxPolA-bearing lineages are marked by stars. For each test tree (Trees 2-9), the difference in log-likelihood from the ML tree (Δlog*L*) and the *P*-value from the AU test are listed.

### Perspective for the mitochondrial DNAP in pre-LECA

It is widely accepted that mitochondrial endosymbiosis occurred before the radiation of all extant eukaryote lineages, and thus, an essentially modern mitochondrion containing a reduced genome relative to its free-living bacterial ancestors was present in the common ancestor of all extant eukaryotes (the last eukaryotic common ancestor or LECA). Considering the α-proteobacterial origin of mitochondria, it is rational to assume that the DNA maintenance had been carried out by the α-proteobacterial DNAPs including Pol`I` in the proto-mitochondrion (a primitive mitochondrion in the pre-LECA stem lineage). Is it then possible that rdxPolA is a direct descendant of the putative proto-mitochondrial Pol`I`? The most recent studies suggested that the proto-mitochondrion represented a lineage phylogenetically sister to essentially all the α-proteobacteria known to science so far (Martijn et al. 2022; Muñoz-Gómez et al. 2022). Intriguingly, our analysis found the intimate affinity of the rdxPolA sequences to, yet outside, the radiation of α-proteobacterial Pol`I` sequences (Fig. 5D), being compatible with the phylogenetic position of the putative bacterial lineage from which the proto-mitochondrion emerged. In the first scenario, multiple secondary losses of this DNAP must have occurred after the divergence of major branches in the tree of eukaryotes and POP supplemented the secondary loss of rdxPolA. Alternatively, the sporadic distribution of rdxPolA in eukaryotes can be explained by lateral gene transfer. This second scenario was not positively supported by the rdxPolA phylogeny, in which the relationship among the discobid, malawimonad, and ancyromonad sequences remains unresolved (Fig. 5D). Overall, we prefer the scenario assuming the vertical inheritance of rdxPolA rather than the other involving eukaryote-to-eukaryote LGT.

To examine whether rdxPolA is derived from the proto-mitochondrion in the LECA, it would be useful to know the type of mitochondrion-localized DNAP in the earliest eukaryotes including LECA, which can in principle be deduced by historical reconstructions guided by the rooted eukaryote phylogeny. Unfortunately, most studies so far carried out to explore the eukaryotic root by using a non-eukaryotic outgroup are by themselves not sufficient to infer the early evolution of mitochondrion-localized DNAP, as they included only Discoba of all the three rdxPolA-containing lineages (Derelle and Lang 2012; He et al. 2014; Al Jewari and Baldauf 2023a; Al Jewari and Baldauf 2023b). However, when the root positions suggested by most of these analyses are mapped onto taxonomically comprehensive unrooted eukaryote phylogenies as recently inferred from multigene datasets (Lax et al. 2018; Tice et al. 2021; Tikhonenkov et al. 2022), the rdxPolA-containing taxa are found in both principal clades defined by the root, with Discoba in one of them the Malawimonadida plus Ancyromonadida in the other. This is consistent with the outgroup-based taxon-rich rooting analysis conducted by Derelle et al. (2015), which splits Discoba and Malawimonadida into the two major clades separated by the inferred root (ancyromonads were missing from the analysis). On the other hand, the analysis by Cerón-Romero et al. (2022), who took a gene tree–species tree reconciliation approach (taking into account gene duplications and losses) to infer the eukaryotic root, does not support rdxPolA as a LECA-resident protein. However, the results reported by these authors need to be interpreted with caution, given the fact that the root position most favoured by their analysis was inside opisthokonts (between fungi and all other eukaryotes), which is at odds with solid evidence for opisthokont monophyly and would require major reconsideration of early eukaryote evolution.

Although the position of the eukaryotic root has yet to be resolved with confidence, the distribution of rdxPolA mapped on the current tree of eukaryotes certainly demands multiple secondary losses of this DNAP. Interestingly, as proposed by previous studies (Moriyama et al. 2011; Hirakawa and Watanabe 2019; Harada and Inagaki 2021) and further reinforced by our expanded sampling, the phylogenetic distribution of POP in eukaryotes implies that the LECA also possessed POP as another mitochondrion-localized DNAP, with secondary losses of POP having occurred in multiple branches of the tree of eukaryotes (e.g., the current rdxPolA-containing lineages). Furthermore, the haptophyte-derived POP genes in choanoflagellates (Fig. S8) demand a careful evaluation of to which degree LGT contributed to the current distribution of POP in eukaryotes. Despite the uncertainties discussed above, in light of the results from our comprehensive survey of famA DNAPs in eukaryotes we propose that both rdxPolA and POP were established in the pre-LECA stage. Such hypothetical ancestral “redundancy” is reminiscent of the history of mitochondrial RNA polymerases (RNAPs), with jakobids possessing the ancestral multisubunit RNAP (encoded by the mitochondrial genome) and all other eukaryotes utilizing a novel nucleus-encode phage-derived single-subunit RNAP (Roger et al. 2017; Gray et al. 2020). Here the co-existence of both RNAPs in the LECA followed by their differential loss also seems to best fit their modern distribution and the phylogenetic position of jakobids among eukaryotes.

## Conclusions

Our analyses uncovered a series of previously unnoticed types of eukaryotic family A DNA polymerases that each occur in multiple representatives of broader protist taxa and trace their origin to evolutionary events at least hundreds of millions of years ago. Two of these DNAP types, not analyzed in detail here (alvPolA and abanPolA), may be novel components of the nuclear machinery of DNA maintenance and deserve specific attention in a separate future study. The remaining eight newly defined DNAP types fundamentally enrich our understanding of the organellar biology. The origin of four of them appears to be directly linked to the emergence of the primary (rgPolA) or different types of secondary (chlnmPolA, eugPolA, cryptoPolA) plastids, and their identification adds so far missing pieces into the mosaic of plastid endosymbioses. Possibly most interesting is the discovery that rdxPolA might represent a previously unknown ancestral (α-proteobacterial) mitochondrial component established pre-LECA and retained by only a minor fraction of extant eukaryote lineages. Further growth of sequence data from eukaryotes, prokaryotes as well as viruses will certainly help to refine our current interpretation of the origin and evolutionary history of the different DNAPs and will resolve open questions such as the nature of the mitochondrial DNAP in Hemimastigophora. Functional genomics approaches need to be employed to define the role of the functionally uncharacterized DNAPs more precisely and to resolve why some mitochondria and plastids seem to possess two different DNAP types. A salient unanswered question is whether organellar genomes may depend on DNAPs from families other than the family A. Except for family B DNAPs serving in the chlorarachniophyte nucleomorph, no such cases have been described, but a systematic study like we have carried out here for the family A is lacking. Clearly, a lot is yet to be learned about how mitochondria and plastids deal with their DNA replication and repair at the paneukaryotic scale.

## Materials and Methods

### Cell cultures and RNA-seq analyses of *Tsukubamonas globosa* and *Fabomonas tropica*

*Tsukubamonas globosa* NIES-1390 was purchased from Microbial Culture Collection at the National Institute for Environmental Studies (https://mcc.nies.go.jp/). *Fabomonas tropica* SRT902 was isolated from seawater sampled at Enoshima Aquarium (Enoshima, Kanagawa, Japan) on October 30^th^, 2019. The species identification of the strain was supported by its 18S rRNA gene being 99% identical to the the respective gene sequence (JQ340336.1) of the type strain *F. tropica* nyk4. The culture of *T. globosa* NIES-1390 was maintained in the laboratory in the 1,000× diluted LB medium containing soil extract and a vitamin mix at 20 °C. The culture of *F. tropica* SRT902 was maintained in the laboratory in seawater containing soil extract and vitamin mix at 20 °C. Soil extract was prepared from soil collected at the Sugadaira Reseach Station, University of Tsukuba (Ueda, Nagano 386-2204, Japan) using a previously described method (Provasoli et al. 1957), and 10 ml was used per liter of cultivating medium. For each liter of cultivating medium, 1 μg of vitamin B_12_, 1 μg of biotin, and 100 μg of thiamine HCl were used as vitamin mix. We harvested the cells from the laboratory cultures of *T. globosa* and then extracted total RNA using the TRIzol reagent and Phasemaker, following the manufacturers’ protocol (MAN0016166). The *T. globosa* RNA sample was subjected to a cDNA library construction using NEBNext Ultra II Directional RNA Library Prep Kit and then to RNA-seq analysis by using the Illumina HiSeq platform at a biotech company (AZENTA Japan Corp., Tokyo, Japan). The experimental procedures for the RNA-seq analysis of *F. tropica* were principally the same, albeit SMART-Seq v4 Ultra Low Input RNA Kit, Nextera DNA Flex Library Prep Kit, and the Illumina HiSeq X platform were applied for cDNA synthesis, cDNA library construction, and sequencing, respectively (the sequencing has been done by Macrogen Japan Corp., Tokyo, Japan). We obtained 64,305,977 and 204,000,750 paired-end reads from the RNA-seq analyses of *T. globosa* and *F. tropica*, respectively. The raw sequence reads were trimmed with fastp v0.23.2 (Chen et al. 2018) with the -q 20 -u 80 option and then assembled with Trinity v2.13.2 (Grabherr et al. 2011) using the default options.

### Phylogeny of famA DNAPs identified in bacteria, phages, and eukaryotes

We searched for novel famA DNAPs using HMMER searches (Eddy 2011) against EukProt3 database (Richter et al. 2022) with a profile HMM generated using HMMER from the seed alignment of the Pfam family DNA_Pol_A (pfam00476, version 34). Significant hits (up to the inclusion threshold) were gathered and preliminary phylogenetic analyses were carried out to identify potentially novel eukaryotic DNAP types. Additional databases were then searched with blastp or tblastn (Camacho et al. 2009) to identify additional members of the putative new types, including the resources provided by GenBank (Sayers et al. 2021), JGI genome portal (https://genome.jgi.doe.gov/portal/), the MMETSP (Keeling et al. 2014) and OneKP transcriptome sequencing projects (Leebens-Mack et al. 2019), and individual sequence datasets deposited in Dryad as supplemental data to publications reporting on the generation of genome or transcriptome assemblies (further details on the sequence sources are provided in Table S1). In addition, we searched the transcriptome assemblies from *T. globosa* and *F. tropica* generated as part of this study, and we also obtained a rdxPolA sequence from a transcriptome assembly from the jakobid *Ophirina amphinema* (the whole dataset to be published elsewhere) (Yabuki et al. 2018). The famA DNAP sequences showed greater than 95% identity to non-eukaryotic sequences by blast analyses against the NCBI non-redundant protein sequence database. We conducted preliminary phylogenetic analyses to identify and discard the DNAP sequences that showed intimate affinities to Polγ, POP, Polθ, Pol`I`A, Pol`I`BCD+, or PREX. The sequences used for further analyses were manually curated to identify and fix possible mistakes in the respective gene models and to evaluate their completeness (especially concerning the N-terminus critical for correct prediction of subcellular localization). In several cases a complete or at least extended sequence was obtained by manually joining partially assembled sequence fragments. We eventually retained 134 sequences as the candidates for previously undescribed famA DNAPs, which are listed with relevant details in Table S1.

For phylogenetic analyses we additionally retrieved a set of reference sequences representing the diversity of Pol`I` in bacteria and phages from the refseq_select_prot database in NCBI as of March 5, 2021 [the detailed procedure was the same as described in Harada and Inagaki (2021)]. The representative sequences of Polγ, POP, Polθ, Pol`I`A, Pol`I`BCD+, and PREX were selected from the alignment generated by some of us before (Harada and Inagaki 2021). The 134 eukaryotic sequences of the candidates for novel famA DNAPs, the Pol`I` sequences of bacteria and phages, and the sequences of the previously known famA DNAP in eukaryotes were aligned by MAFFT v7.490 with the L-INS-i model (Katoh and Standley 2013). The aa sequences encoding the polymerase domain is the only part of the protein that is shared among all the sequences analyzed. Further, ambiguously aligned positions were excluded manually and gap-containing positions were trimmed by trimAl v1.4 (Capella-Gutiérrez et al. 2009) from the initial alignment, leaving 355 unambiguously aligned aa positions in 488 famA DNAPs (“global DNAP” alignment). We then subjected the global DNAP alignment to the ML tree reconstruction by using IQ-TREE v.2.2.0 with the LG+C60+F+R10 model (Minh et al. 2020) that was selected by ModelFinder with “-mset LG+C60” option (Kalyaanamoorthy et al. 2017). Statistical support for each bipartition in the ML tree was calculated by 1000-replicate an ultrafast bootstrap approximation (Hoang et al. 2018).

### Phylogenetic analyses assessing the origins of novel types of famA DNAPs in eukaryotes

The origins of the ten novel types of famA DNAPs in eukaryotes—acPolA, rgPolA, chlnmPolA, eugPolA, pyramiPolA, chloroPolA, rdxPolA, cryptoPolA, abanPolA, and alvPolA—were assessed by separate phylogenetic analyses. Six of the DNAPs (acPolA, rgPolA, chlnmPolA, chloroPolA, abanPolA, and alvPolA) were each analyzed as part of a separate alignment, whereas the other four were combined in pairs (eugPolA plus pyramiPolA, rdxPolA plus cryptoPolA) into two separate alignments based on the fact the DNAPs in pairs were closely related in the global DNAP phylogeny. Each of the eight alignments was supplemented by a set of bacterial sequences selected by targeted searches to include the putative closest relatives of the eukaryotic DNAPs analyzed. Specifically, the bacterial Pol`I` sequences were retrieved by running a BLASTP search against the GenBank Refseq_selec_prot database with the eukaryotic famA DNAP sequences of interest as queries. We kept approximately the top 1,000 bacterial Pol`I` sequences matched to the queries in each of the BLAST surveys. The redundancy in the set of bacterial Pol`I` sequences retrieved from the database was removed by using CD-HIT (Li and Godzik 2006). The *E*-value cutoffs for the BLAST surveys and parameters for CD-HIT are listed in Table S2. The query sequences (one or two types of the novel famA DNAPs in eukaryotes) and the bacterial Pol`I` sequences retrieved from the corresponding BLAST survey were aligned by using MAFFT with the L-INS-i model. We then polished the eight alignments by manual trimming of ambiguously aligned positions and exclusion of gap-containing positions by trimAl. As a result, the alignments built for acPolA, chlnmPolA, chloroPolA, abanPolA, and alvPolA plus the respective sets of bacterial Pol`I` sequences were comprised of unambiguously aligned positions in the polymerase domain only, whereas the final alignments made for rgPolA, eugPolA/pyramiPolA, and rdxPolA/cryptoPolA plus their bacterial homologs were longer for including also unambiguously aligned positions in the 5′-3′ exonuclease and the 3′-5′ exonuclease domains. The eight alignments were subjected to ML tree search and 100-replicate nonparametric ML bootstrap analysis by using IQ-TREE. The substitution models applied for the tree searches were selected by ModelFinder. For the nonparametric ML bootstrap analysis, we applied the same model as the corresponding tree search but incorporated PMSF (posterior mean site frequencies) that were calculated over the ML tree as the guide (Wang et al. 2018). The eight alignments were also subjected to Bayesian analysis with the CAT+GTR+Γ model using PhyloBayes v1.8 (Lartillot et al. 2013). For each alignment, two Markov chain Monte Carlo runs were run to calculate the consensus tree and Bayesian posterior probabilities. Details on the eight alignments (i.e., the numbers of the DNAP sequences and positions used for the phylogenetic analyses) and the substitution models selected for the phylogenetic analyses are provided in Table S2.

### *In silico* prediction of subcellular localization and functional domains of famA DNAPs

Seventy-five out of the 134 sequences representing the novel famA DNAP types identified in this study were considered to have complete and accurately defined N-termini. We subjected their N-terminal regions to *in silico* prediction of the targeting potential to subcellular compartments (i.e., endoplasmic reticulum, mitochondria, and plastids) using a suite of tool, namely MitoFates (Fukasawa et al. 2015), NommPred (Kume et al. 2018), DeepLoc-1.0 (Almagro Armenteros et al. 2017), DeepLoc-2.0 (Thumuluri et al. 2022), TargetP-1.1 (Emanuelsson et al. 2000), TargetP-2.0 (Almagro Armenteros et al. 2019), PredSL (Petsalaki et al. 2006), and TMHMM-2.0 (Krogh et al. 2001). Additionally, we subjected the full-length pyramiPolA and chloroPolA sequences to PredAlgo (Tardif et al. 2012), which was developed for predicting the subcellular localization of green algal proteins. Conserved functional domains in the 134 famA DNAPs were predicted using InterProScan v5.55 with the Pfam database (Jones et al. 2014; Paysan-Lafosse et al. 2023).

### Experiments assessing the subcellular localizations of the selected famA DNAPs

#### rdxPolAs

The N-terminal amino acid sequences predicted as mitochondrial targeting signals were fused to green fluorescent protein (GFP), and the fusion proteins were expressed in yeast (*Saccharomyces cerevisiae*). In the five rdxPolA sequences of *Ophirina amphinema*, *Gefionella okellyi*, *Naegleria gruberi*, *F. tropica*, and *T. globosa*, the first 44, 26, 95, 71, and 76 amino acid residues were predicted as MTS by MitoFates, respectively. Double-stranded DNA (dsDNA) fragments corresponding to regions encoding the putative MTS aa sequences of the five rdxPolA proteins were synthesized with two modifications described below. First, the codon usage, which may be specific to the five eukaryotic genomes, was optimized for expression in yeast. Second, the synthesized nucleotide sequences were flanked with the sequence recognized by *Eco*RI (5′-GAATTC-3′) and that by *Bam*HI (5′-GGATCC-3′) at the 5′ and 3′ termini, respectively. The synthesized dsDNA fragments were inserted into pYX142-mtGFP vector (Westermann and Neupert 2000) via the restriction sites mentioned above. The dsDNA synthesis and vector construction were done by a biotech company (AZENTA Japan Corp, Japan). One μg of the vectors was transformed in the BY4742 strain of yeast (Brachmann et al. 1998). Yeast transformations were performed according to the lithium acetate procedure (Schiestl and Gietz 1989), and transformants were selected on a synthetic dextrose minimal medium lacking lysine, histidine, and uracil. Eight to 30 days after transformation, mitochondria in yeast transformants were stained with MitoTracker Red at a final concentration of 0.5 μM and were observed with an Olympus BX51 fluorescent microscope (Olympus) equipped with an ORCA-3CCD color camera (Hamamatsu Photonics).

#### *Bigelowiella natans* chlnmPolA

To confirm the subcellular localization of the *B. natans* chlnmPolA, a GFP-tagged protein was heterologously expressed in the chlorarachniophyte *Amorphochlora amoebiformis*. The *B. natans* chlnmPolA was predicted to have an N-terminal extension containing a signal peptide (the first 24 amino acid residues) by TargetP 2.0 (Almagro Armenteros et al. 2019). We constructed a plasmid vector expressing GFP fused with the first 100 aa residues of *B. natans* chlnmPolA. Firstly, total RNA was extracted from *B. natans* CCMP621 cells using TRIzol reagent (Invitrogen), and cDNA was synthesized using SuperScript IV Reverse Transcriptase (Invitrogen) with an oligo(dT) primer. Secondly, the cDNA fragment encoding chlnmPolA was amplified by PCR with a pair of specific primers (5′-ATGATCAGGAAGCAATATATGTTGAG-3′ and 5′-CACCAGAGAGTTAGATCCCAT-3′), and inserted at the 5’ of the GFP gene in the plasmid vector pLaRGfp (Hirakawa et al. 2009) using GeneArt Seamless Cloning and Assembly Enzyme Mix (Invitrogen). The plasmid was introduced into *Escherichia coli* DH5α, and the inserted sequence was verified by Sanger sequencing. To express the GFP fusion protein, *A. amoebiformis* CCMP2058 cells were transformed with 10 μg of the plasmid using a Gene Pulser Xcell electroporation system (Bio-Rad), as described previously (Fukuda et al. 2020). One to two days after transformation, GFP fluorescence was observed under an inverted Zeiss LSM 510 laser scanning microscope (Carl Zeiss).

#### *Guillardia theta* cryptoPolA and POPs

To confirm the subcellular localization of the *G. theta* cryptoPolA and the two POPs, the N-terminal amino acid sequences of the three DNAPs fused to GFP were heterologously expressed in the diatom *Phaeodactylum tricornutum* UTEX642. For the three DNAPs, dsDNA fragments of the region from the start codon until the beginning of the first functional domain were synthesized by (AZENTA Japan Corp, Japan). Each of the synthesized DNA fragments was amplified by PCR and inserted at the 5’ of the GFP gene in the pPha-NR vector (NovoPro Bioscience) using NEBuilder HiFi DNA Assembly (New England BioLabs). Five μg of the pPha-NR constructs were introduced into *P. tricornutum* using a NEPA21 gene gun (NEPAGENE) (Miyahara et al. 2013) and grown the transformants in a zeocin-based selection medium for 57 to 133 days. Actively growing *P. tricornutum* transformants in a zeocin-based selection medium were observed with an Olympus BX51 fluorescent microscope (Olympus) equipped with an Olympus DP72 CCD color camera (Olympus). Mitochondria of transformants transfected with fragments of POP1 were stained with MitoTracker Orange at a final concentration of 0.1 μM.

### Phylogenomic analysis of eukaryotes

We updated the 351-gene phylogenomic alignment used for Yazaki et al. (2022). The amino acid sequences included in each of the 351 single-gene alignments generated in the previous study and the homologous sequences of *T. globosa* and *F. tropica* SRT902, which were retrieved from the RNA-seq data, were combined and re-aligned. After the exclusion of ambiguously aligned positions, we subjected the 351 updated alignments separately to preliminary ML phylogenetic analyses. Among the 351 alignments, we omitted 10 alignments containing putative paralogous sequences, and a single alignment, in which the sequence sampling from ancyromonads were poor, from the final ML phylogenetic inferences. The remaining 340 updated alignments were concatenated into a single “phylogenomic” alignment comprising 97 taxa and 116,499 unambiguously aligned amino acid positions. We analyzed the 340-gene phylogenomic alignment with the ML method by using IQ-TREE (Minh et al. 2020) with the LG+C60+F+Γ model. The statistical support for each bipartition was calculated by 100-replicate nonparametric ML bootstrap analysis. For the bootstrap analysis, we applied the LG+C60+F+Γ+PMSF model (the ML tree inferred with the LG+C60+F+Γ model was used as the guide).

We tested the possibility that the rdxPolA-bearing lineages form a monophylum (synonymous with the monophyly of the POP-bearing assemblages) using an AU test (Shimodaira 2002). Six alternative trees were generated from the ML tree (see above) by pruning and regrafting the Amorphea plus CRuMs clade from the original position to six alternative positions (Trees 2-7; Fig. 7B), followed by the re-optimization of the internal branching pattern in each subclade. We additionally inferred the best trees under two distinct topological constraints, one enforcing the Amorphea plus CRuMs clade and Diaphoretickes tied together, and the other enforcing the clade of all the rdxPolA-bearing lineages (Trees 8 and 9; Fig. 7B). Note that trees 7 and 9 were identical to each other, except the internal relationship among stramenopiles (see supplementary materials for the details). We calculated the site-wise log-likelihoods (site-lnLs) over the ML and eight alternative trees (the substitution model used for the ML tree search was applied). The resultant site-lnL data were then subjected to an AU test. IQ-TREE (Minh et al. 2020) was used for all the calculations required for the test described above.

## Data availability

All the alignments, treefiles inferred from the alignments, and the fasta files containing transcriptome assemblies generated in this study are available at https://doi.org/10.5061/dryad.b5mkkwhjz. The transcriptome data of *T. globose* NIES-1390 and *F. tropica* SRT902 were deposited in GenBank/EMBL/DDBJ Sequence Read Archive under accession no. PRJNA932580.

## Acknowledgments

We would like to thank the Enoshima Aquarium (Kanagawa, Japan) for providing the seawater samples and Dr. Takashi Shiratori (University of Tsukuba) for establishing the *F. tropica* strain. The work was supported by the Japan Society for Promotion of Sciences projects 18KK0203, 19H03280, 23H02535, and BPI05044 (to Y.I.), 22J11104 (to R.H.), 19H03274 (to R.K.), and the Czech Science Foundation grant no. 21-19664S (to M.E.). *T. globosa* (NIES-1390) and *S. cerevisiae* BY4742 strain were provided by the NIES and YGRC through the National Bio-Resource Project (NBRP) of the Ministry of Education, Culture, Sports, Science and Technology (MEXT), Japan, respectively. Computations were partially performed on the NIG supercomputer at ROIS National Institute of Genetics. The phylogenetic analyses conducted in this work have been carried out partially under the ‘Interdisciplinary Computational Science Program’ in the Center for Computational Sciences, University of Tsukuba.

