## Supplementary material for "Encyclopaedia of family A DNA polymerases localized in organelles: Evolutionary contribution of bacteria including the proto-mitochondrion": Fig. S1-S8

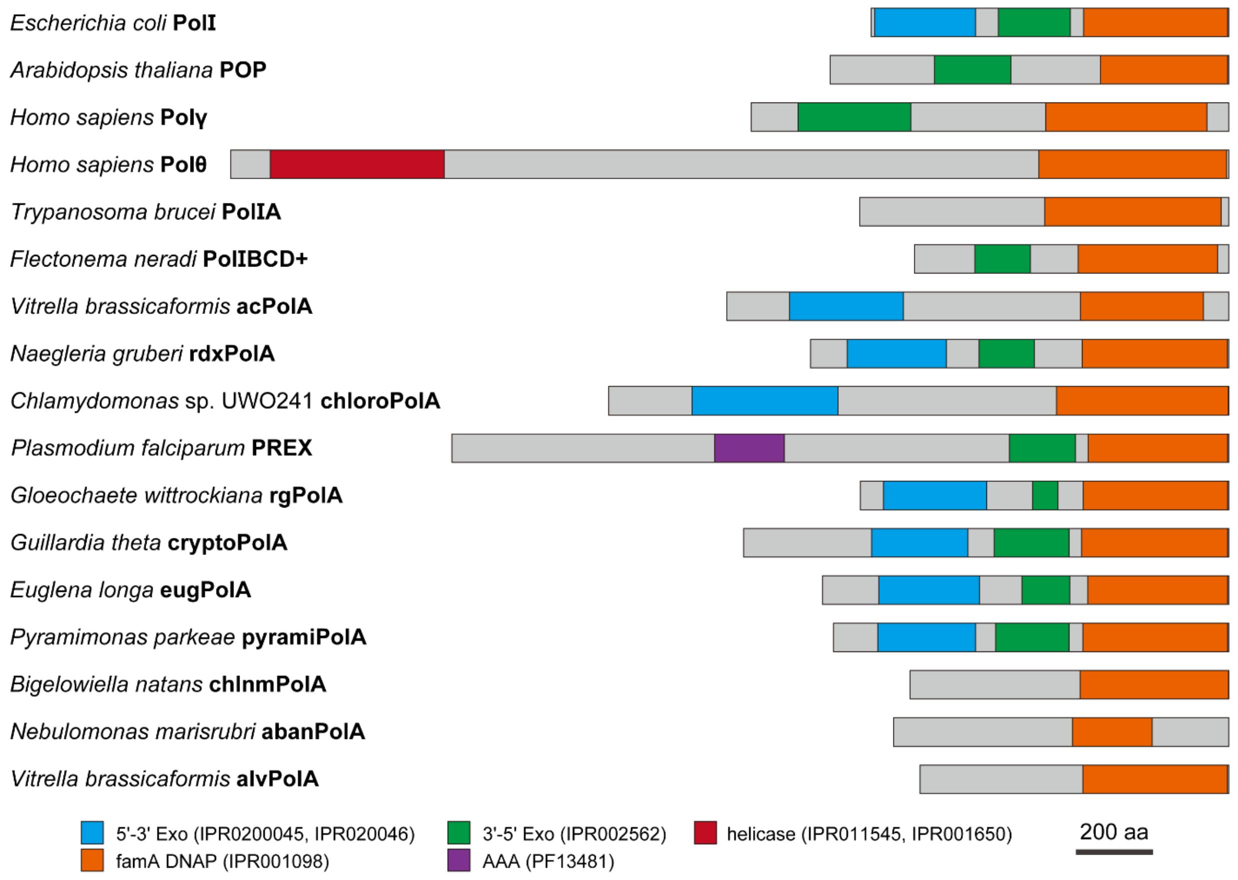


Fig. S1. Domain structures of DNA polymerase I (PolI) of *Escherichia coli* and PolI-related (family A) DNA polymerases in eukaryotes. Each type is represented by a single selected member. The 3'-5' exonuclease domains (IPR0200045 and IPR020046), 5'-3' exonuclease domain (IPR002563), family A DNA polymerase domain (IPR001098), AAA ATPase domain (PF13481), and helicase domain (IPR011545, IPR001650) are shown in blue, green, orange, purple, and red, respectively.

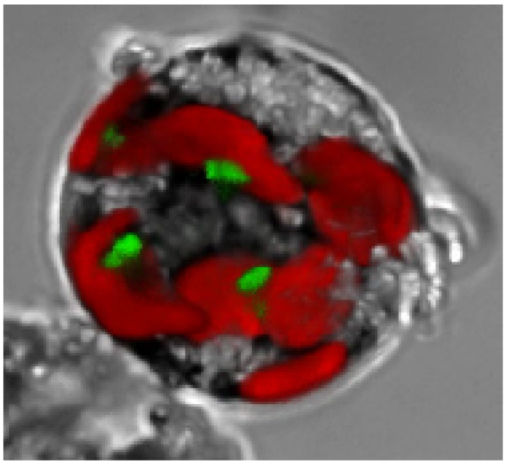

Fig. S2. The maximum intensity projection from a Z-stack of an *Amorphochlora amoebiformis* cell. The cell expressed the green fluorescent protein (GFP) fused with the N-terminus of the chlnmPolA protein of *Bigelowiella natans*. Red, chlorophyll-autofluorescence; green, GFP.

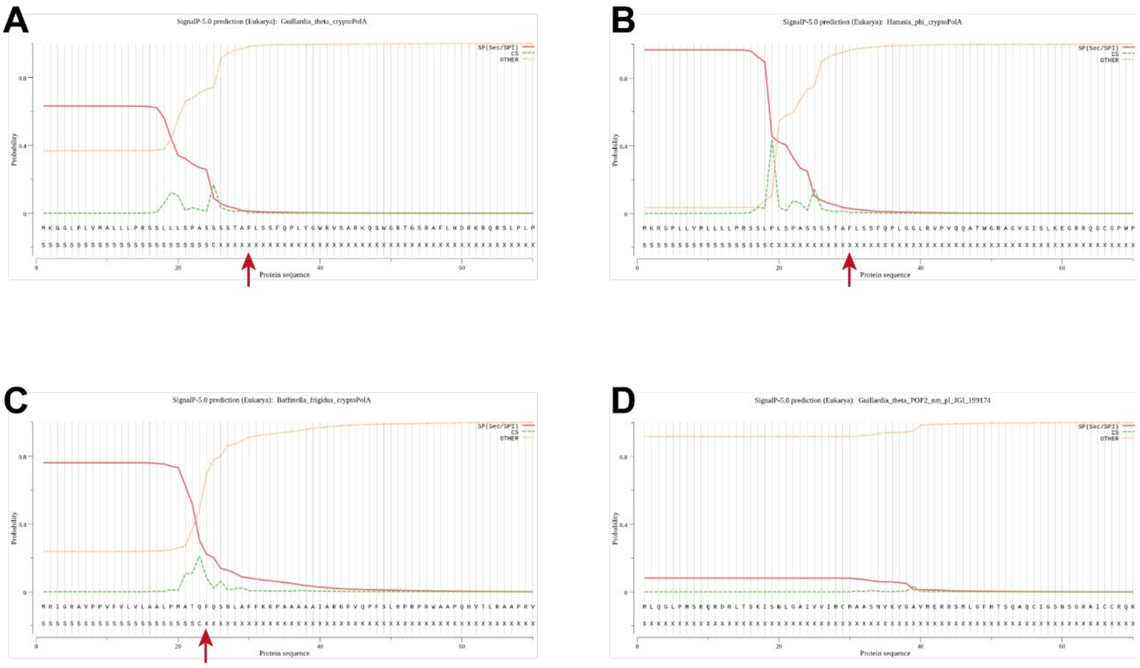


Fig. S3. Prediction of a signal peptide in the N-termini of selected famA DNAPs identified in cryptophytes. We used SignalP-5.0 for the predictions. (A) *Guillardia theta* cryptoPolA, (B) *Hanusia phi* cryptoPolA, (C) *Baffinella frigidus* cryptoPolA, and (D) *Guillardia theta* POP2. The phenylalanine residues closest to the predicted cleavage sites in the three cryptoPolA sequences are highlighted with red arrows.

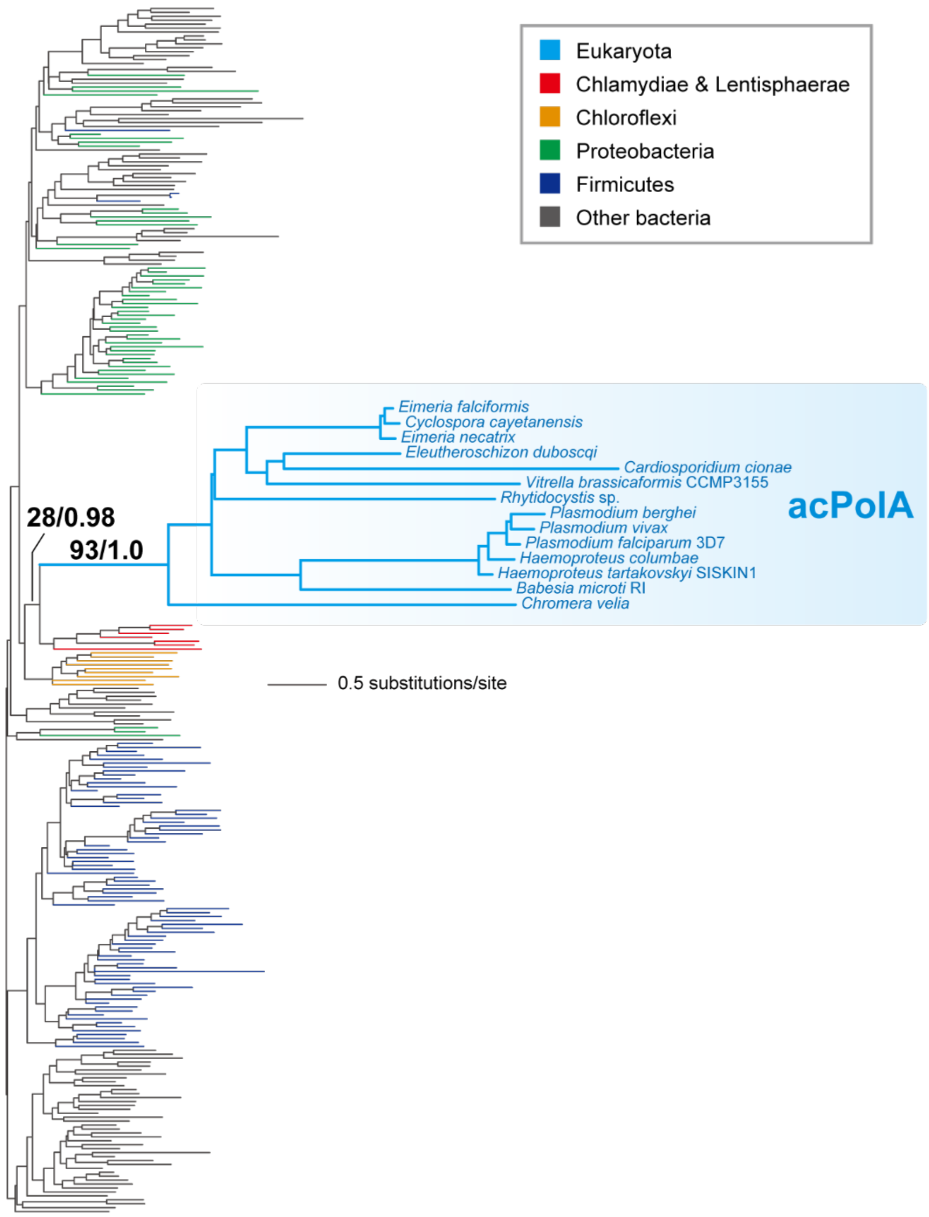


Fig. S4. Maximum-likelihood (ML) phylogeny assessing the origin of acPolA. The alignment comprised 263 DNAP sequences with 295 unambiguously aligned positions. The ML nonparametric bootstrap support values and Bayesian posterior probabilities (if they are equal to or greater than 0.5) are presented only for the key nodes for the origin of acPolA. The clades/branches are color-coded, as shown in the inset.

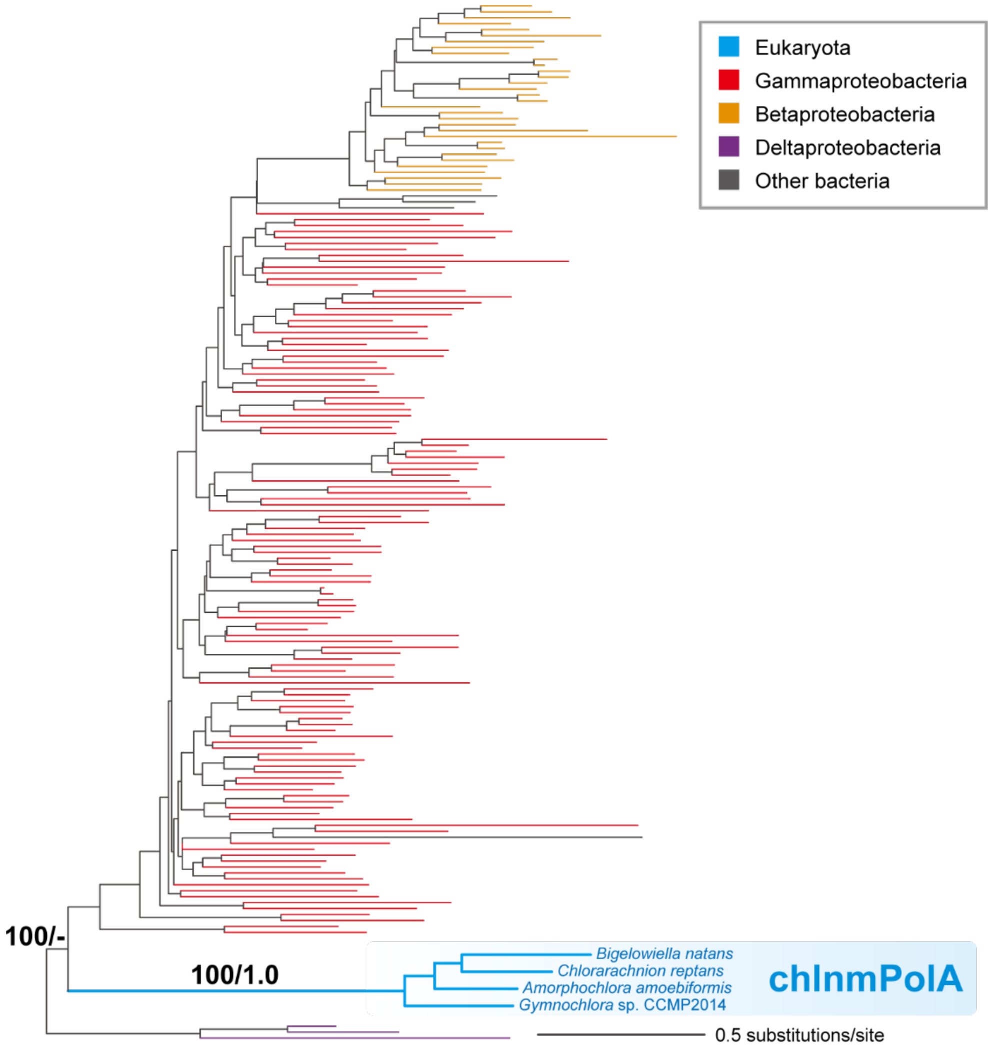


Fig. S5. Maximum-likelihood (ML) phylogeny assessing the origin of chlnmPolA. The alignment comprised 164 DNAP sequences with 392 unambiguously aligned positions., The ML nonparametric bootstrap support values and Bayesian posterior probabilities (if they are equal to or greater than 0.5) are presented only for the key nodes for the origin of chlnmPolA. The clades/branches are color-coded, as shown in the inset.

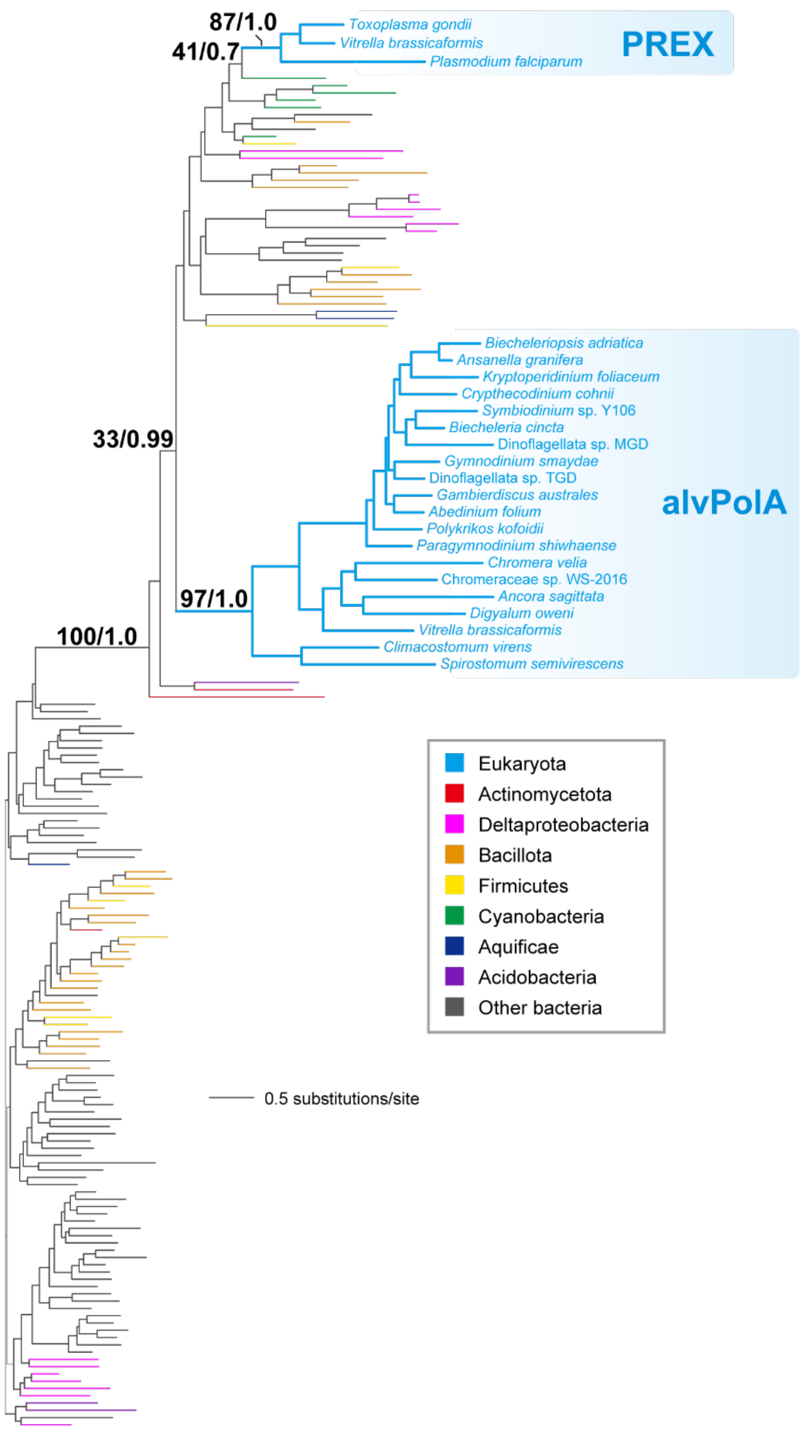


Fig. S6. Maximum-likelihood (ML) phylogeny assessing the origin of alvPolA. The alignment comprised 161 DNAP sequences with 388 unambiguously aligned positions. The ML nonparametric bootstrap support values and Bayesian posterior probabilities (if they are equal to or greater than 0.5) are presented only for the key nodes for the origin of alvPolA. The clades/branches are color-coded, as shown in the inset.

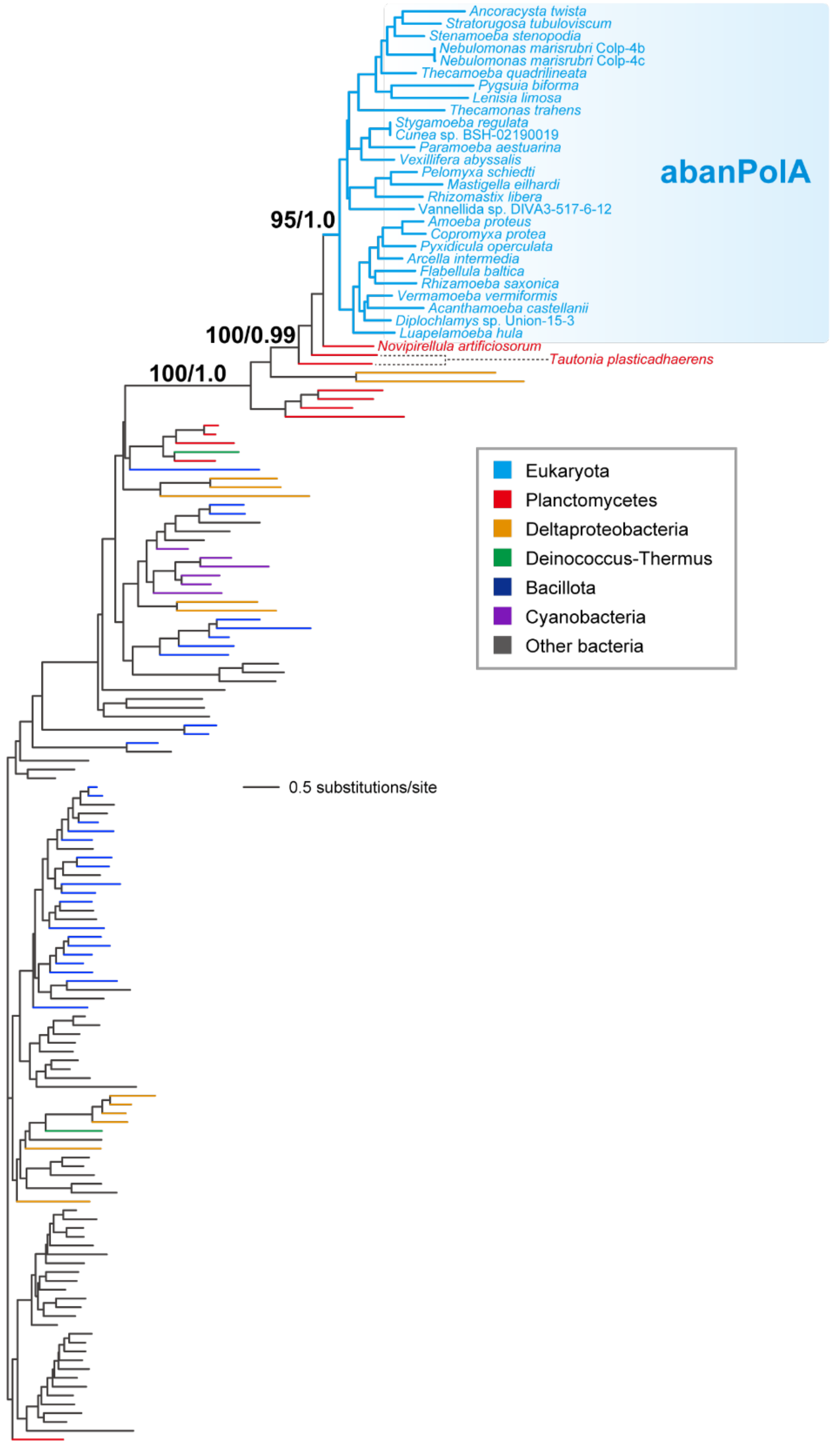


Fig. S7. Maximum-likelihood (ML) phylogeny assessing the origin of abanPolA. The alignment comprised 152 DNAP sequences with 380 unambiguously aligned positions. The ML nonparametric bootstrap support values and Bayesian posterior probabilities (if they are equal to or greater than 0.5) are presented only for the key nodes for the origin of abanPolA. The clades/branches are color-coded, as shown in the inset.

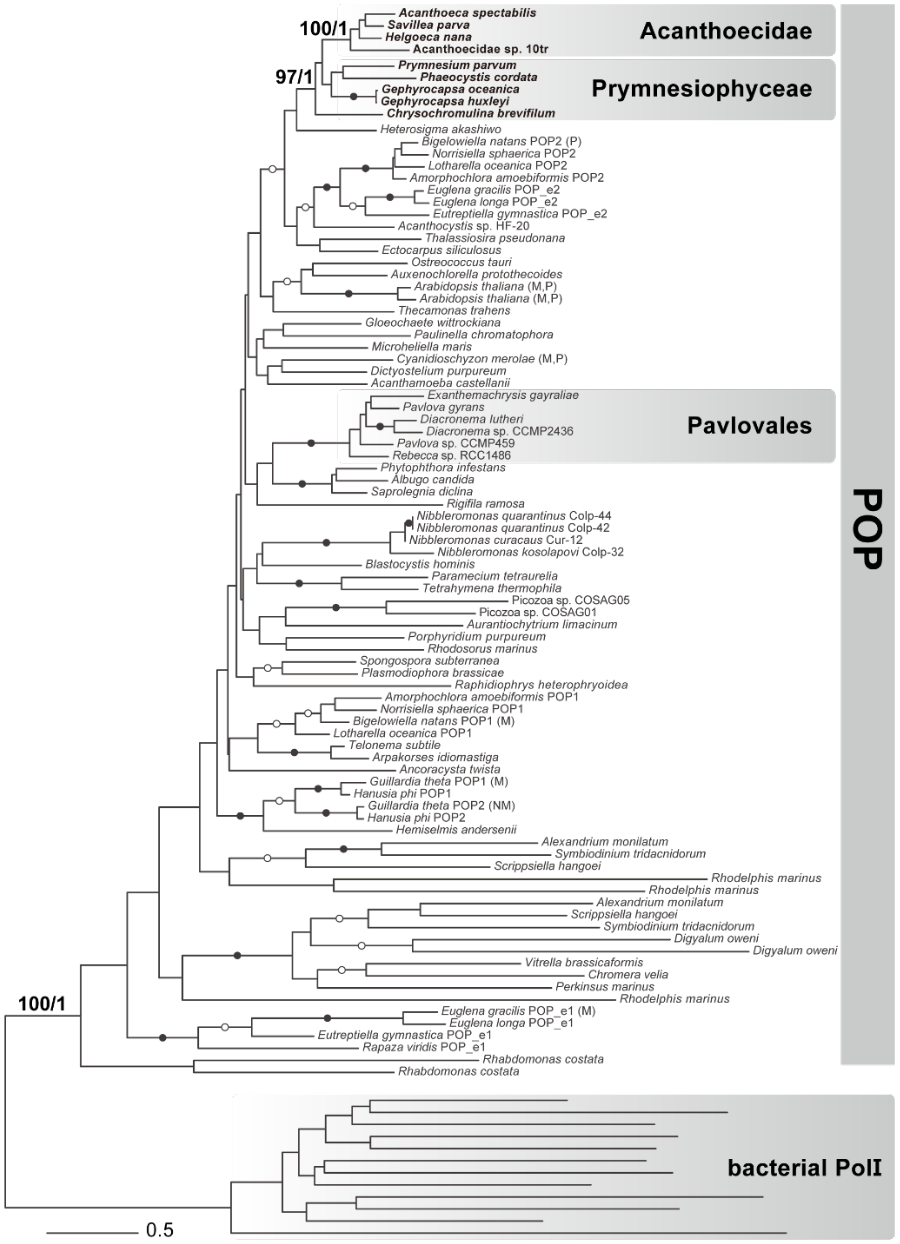


Fig. S8. Maximum-likelihood (ML) phylogeny of POP. The alignment comprised 88 POP and 12 bacterial PolI sequences with 365 unambiguously aligned positions. The ML nonparametric bootstrap support values (MLBPs) and Bayesian posterior probabilities (BPPs) are presented only for the key nodes for the origin of Acanthoecidae POP. The nodes supported by both MLBPs of 95-100% and BPPs of 1.0 are marked with closed dots. Likewise, the nodes supported by both MLBPs of 70-94% and BPPs of 0.95-0.99 are marked with open dots. Sequences with experimentally confirmed subcellular localization are indicated by localization in parentheses. The POP sequences found in this study, those analyzed in Harada et al. (2020), and the DNAP sequences that represent the diversity of bacterial PolI were aligned by using MAFFT with the L-INS-i model. Ambiguously aligned positions were trimmed manually, and gap-containing positions were trimmed by trimAl. The details of the phylogenetic analyses of the POP alignment are summarized in Table S2.
