## Supplementary material for "Encyclopaedia of family A DNA polymerases localized in organelles: Evolutionary contribution of bacteria including the proto-mitochondrion": Table S2

TABLE S2. Parameters set for the phylogenetic analyses conducted in this study.

|  | BLASTP | CD-HIT | trimAl | Alignment | | IQ-TREE | PhyloBayes | | |
| --- | --- | --- | --- | --- | --- | --- | --- | --- | --- |
|  | e-value | -c | -gt | OTU | aa residue | model | cycles | burn-in | maxdiff |
| global FamA (Fig. 1) | - | - | 0.9 | 488 | 355 | LG+C60+F+R10 | - | - | - |
| rgPolA (Fig. 5A) | 1e-110 | 0.6 | 0.97 | 147 | 781 | LG+C60+F+R9 | 100000 | 10000 | 0.2253 |
| chloroPolA (Fig. 5B) | 1e-100 | 0.6 | 0.9 | 192 | 401 | LG+C60+F+R10 | 100000 | 10000 | 0.2691 |
| eugPolA/pyramiPolA (Fig. 5C) | 1e-130 | 0.65 | 0.9 | 161 | 893 | LG+C60+F+R10 | 100000 | 10000 | 0.2469 |
| rdxPolA/cryptoPolA (Fig. 5D) | 1e-130 | 0.7 | 0.95 | 229 | 782 | LG+C60+F+R10 | 150000 | 70000 | 0.1962 |
| acPolA (Fig. S4) | 1e-50 | 0.6 | 0.8 | 263 | 295 | LG+C60+R10 | 100000 | 20000 | 0.2947 |
| chlnmPolA (Fig. S5) | 1e-125 | 0.7 | 0.99 | 164 | 392 | LG+C60+R10 | 100000 | 10000 | 0.3396 |
| alvPolA/PREX (Fig. S6) | 1e-45 | 0.55 | 0.9 | 161 | 388 | LG+C60+F+R9 | 100000 | 20000 | 0.1147 |
| abanPolA (Fig. S7) | 1e-11 | 0.6 | 0.9 | 152 | 380 | LG+C60+F+I+R9 | 100000 | 10000 | 0.0640 |
| POP (Fig. S8) | - | - | 0.8 | 100 | 365 | LG+C60+F+R6 | 100000 | 10000 | 0.1697 |

­
